# An awareness-dependent saliency map in the human visual system

**DOI:** 10.1101/2021.01.07.424631

**Authors:** Lijuan Wang, Ling Huang, Mengsha Li, Xiaotong Wang, Shiyu Wang, Yuefa Lin, Xilin Zhang

**Affiliations:** Key Laboratory of Brain, Cognition and Education Sciences (South China Normal University), Ministry of Education, Guangzhou, Guangdong 510631, China; School of Psychology, South China Normal University, Guangzhou, Guangdong 510631, China; Center for Studies of Psychological Application, South China Normal University, Guangzhou, Guangdong 510631, China; Guangdong Provincial Key Laboratory of Mental Health and Cognitive Science, South China Normal University, Guangzhou, Guangdong 510631, China

**Keywords:** saliency map, awareness, gradient manner, winner-take-all manner, fMRI

## Abstract

The bottom-up contribution to the allocation of exogenous attention is a saliency map. However, how the saliency map is distributed when multiple salient stimuli are presented simultaneously and how this distribution interacts with awareness remain unclear. These questions were addressed here using visible and invisible stimuli that consisting of two salient foregrounds: the high one served as the target and the low one served as the distractor, which did or did not interfere the target’ saliency, indicating a gradient or winner-take-all manner, respectively. By combining psychophysics, fMRI, and effective connectivity analysis, we found that the saliency map was distributed as a gradient or winner-take-all manner with and without awareness, respectively. Crucially, we further revealed that the gradient manner was derived by feedback from pIPS, whereas the winner-take-all manner was constructed in V1. Together, our findings indicate an awareness-dependent saliency map and reconcile previous, seemingly contradictory findings on the saliency map.

## Introduction

Attentional selection is the mechanism by which a subset of incoming information is processed preferentially. Numerous studies have demonstrated that this attentional selection can either be executed voluntarily by top-down signals derived from goals, such as when directing gaze to an interesting book (Baluch and Itti, 2011; Corbetta and Shulman, 2002; Kanwisher and Wojciulik, 2000; Kastner and Ungerleider, 2000; Serences and Yantis, 2006; Zhang et al., 2016, 2018) or automatically by bottom-up signals from salient stimuli, such as a vertical bar among horizontal bars (Corbetta and Shulman, 2002; Hegdé and Felleman, 2003; Jonides, 1981; Koch and Ullman, 1985; Nakayama and Mackeben, 1989). Throughout this study, we use the term salience to refer to this bottom-up attraction of attentional selection. While a saliency map is defined as a topographical map to describe and predict the distribution of this bottom-up attraction based on a visual input (Itti and Koch, 2001; Koch and Ullman, 1985). Although the bottom-up attraction is typically quick and potent (Jonides, 1981; Nakayama and Mackeben, 1989), little is known about how this bottom-up attraction will be distributed when multiple salient stimuli are presented simultaneously and instantaneously.

A dominant model of the saliency map developed by Itti and Koch (2001) presumed that the bottom-up attention is sequentially allocated, in a winner-take-all manner, to the most salient location from multiple salient regions, which was then suppressed by the inhibition-of-return mechanism (Klein, 2000) so that the bottom-up attention can focus onto the next most salient location (Koch and Ullman, 1985; Wolfe, 1994), and repeating this process generates attentional scanpaths in the stimulus. Accordingly, when a stimulus contains two salient regions with different levels of saliency (for example, 25° and 90° orientation contrasts, *Figure 1A*); the bottom-up attention will be captured by the more salient (i.e., the 90° orientation contrast) region at any given moment in time. To date, however, this model lacks empirical supports since most of previous studies used the stimulus with a single salient region only (Burrows and Moore, 2009; Buschman and Miller, 2007; Chen et al., 2016; Geng and Mangun, 2009; Katsuki and Constantinidis, 2012; Serences and Yantis, 2007, but see Bogler et al., 2011, White et al., 2017a). It is still unclear whether the bottom-up attention is completely allocated to the most salient region (i.e., the winner-take-all manner) or proportionally allocated to all salient regions based on their different levels of saliency (i.e., the gradient manner).

**Figure 1.**
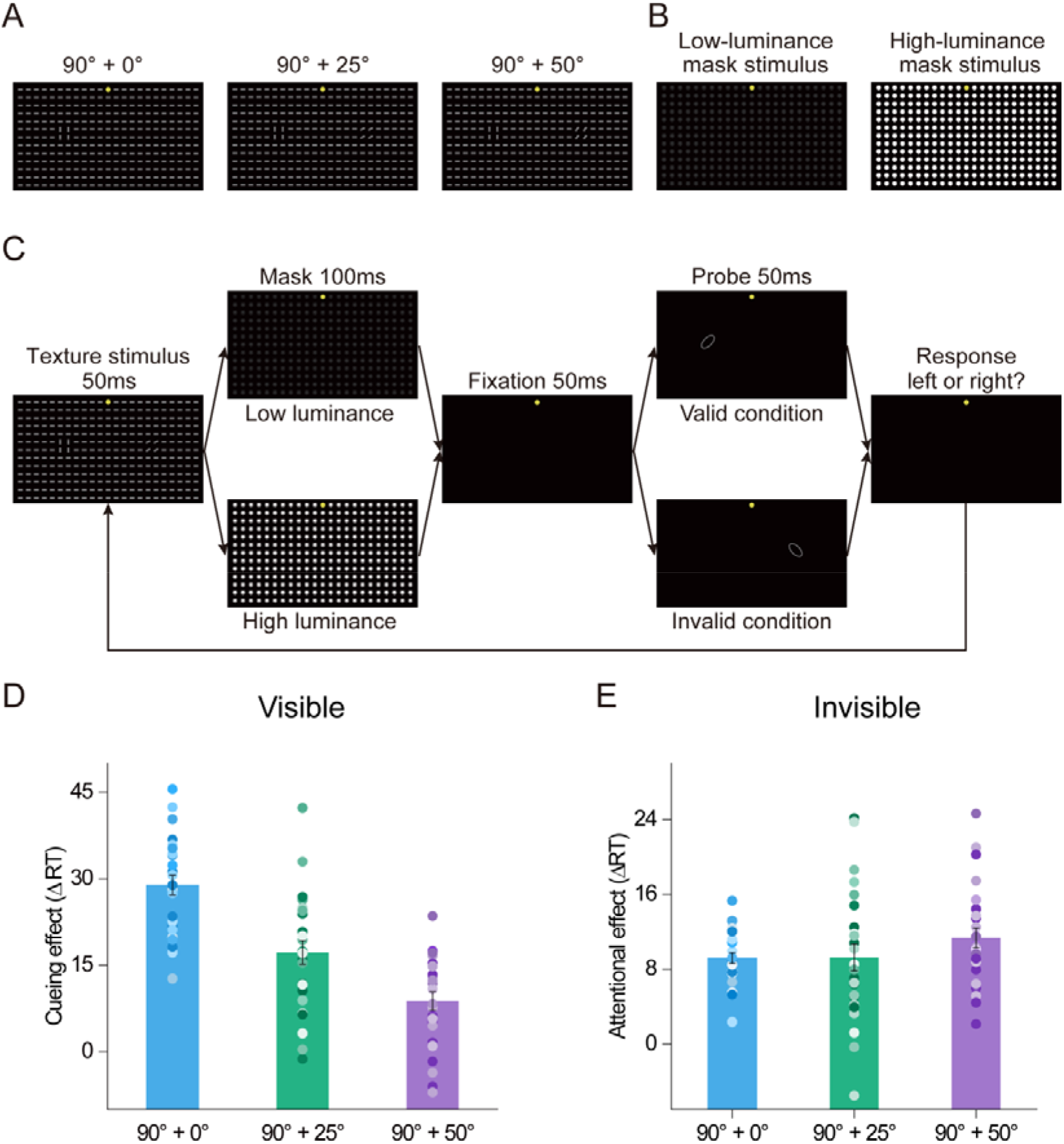
Stimuli, Psychophysical Protocol and Data. (**A**) Three low-luminance texture stimuli presented in the lower visual field (left: 90° + 0°; middle: 90° + 25°; right: 90° + 50°, and the yellow dots indicate the fixation point). Each texture stimulus contained a pair of salient foregrounds centered in the lower left and lower right quadrants (in this case, the 90°-foreground in the lower left quadrant), the high salient one served as the target and the low salient one served as the distractor. (**B**) Low- (left) and high-luminance (right) mask stimuli used in the Visible and Invisible conditions, respectively. (**C**) Psychophysical protocol. A texture stimulus was presented for 50 ms, followed by a 100-ms mask and another 50-ms fixation interval. Then an ellipse probe was presented at randomly either the target location (e.g., the 90°-foreground location, valid cue condition) or its contralateral counterpart (the distractor location, i.e., invalid cue condition) with equal probability. The ellipse probe was orientated at 45° or 135° away from the vertical. Subjects were asked to press one of two buttons as rapidly and correctly as possible to indicate the orientation of the ellipse probe (45° or 135°). (**D** and **E**) The cueing effect of the 90°-foreground (i.e., the target) for 90° + 0°, 90° + 25°, and 90° + 50° texture stimuli in Visible (**D**) and Invisible (**E**) conditions. Each cueing effect was quantified as the difference between the reaction time of the probe task performance in the invalid cue condition and that in the valid cue condition. Error bars denote 1 SEM calculated across subjects and colored dots denote the data from each subject.

In addition, there has been a long-standing debate about the neural loci of the saliency map. Evidence from numerous neurophysiological and imaging studies have shown that the superior colliculus (SC, Fecteau and Munoz, 2006; Kustov and Robinson, 1996; White et al., 2017a; 2017b), pulvinar (Shipp, 2004), substantia nigra (Basso and Wurtz 2002), parietal cortex (Bisley and Goldberg, 2010; Bogler et al., 2011; Buschman and Miller, 2007; Geng and Mangun, 2009; Gottlieb et al., 1998; Serences et al., 2005), V4 (Burrows and Moore, 2009; Mazer and Gallant, 2003), ventral attention network (Asplund et al. 2010; Corbetta and Shulman, 2002), frontal eye fields (Bogler et al., 2011; Serences and Yantis, 2007; Thompson and Bichot, 2005), and the dorsolateral prefrontal cortex (Katsuki and Constantinidis, 2012; Squire et al., 2013) could realize the saliency map. Most of these studies were consistent with the dominant view by Itti and Koch (2001), which proposes that saliency results from pooling different visual features, being independent of whether the feature distinction making a location salient is in color, orientation, or other features (Koch and Ullman, 1985; Wolfe, 1994). Accordingly, higher cortical areas, particularly the parietal and frontal cortex, whose neurons are less selective to specific visual features, are more likely to be possible candidates that realize the saliency map. By contrast, Li (1999, 2002) proposed that primary visual cortex (V1) creates the saliency map via intra-cortical interactions that are manifest in contextual influences (Allman et al., 1985). The saliency of a location is monotonically related to the highest neural response among all the V1 cells that cover that location with their spatial receptive fields (relative to the V1 responses to the other locations), regardless of the preferred feature of the most responsive neuron. This theory has also been supported by several psychophysical (Koene and Zhaoping, 2007; Zhaoping, 2008; Zhaoping and May, 2007; Zhaoping and Zhe, 2015), neurophysiological (Kastner et al., 1997; Nothdurft et al., 1999; Yan et al., 2018), and brain imaging (Chen et al., 2016; Zhang et al., 2012) studies.

An important reason of this controversy is that most of the observed neural substrates for the saliency map are also involved in top-down attentional selection, so that the saliency map is easily and inadvertently contaminated with the top-down signals, such as feature perception, object recognition, and subjects’ intentions (Zhang et al., 2012). Here, to investigate how the bottom-up attraction will be distributed among multiple salient regions, on the one hand, it is thus important to probe bottom-up attractions free from top-down influences. One way to obviate this is to use the backward masking paradigm in which the salient stimuli are presented so briefly and followed by a high luminance mask that they are invisible to subjects. On the other hand, the salient stimulus, visible versus invisible, offers a unique opportunity to reveal how its saliency map interacts with awareness.

As such stimuli, we used both visible (Experiment 1) and invisible (Experiment 2) textures made from bars (Figure 1A). Each texture stimulus contained two salient foregrounds whose bars were oriented differently from the bars in the otherwise uniform background, the high one served as the target and the low one served as the distractor, which did or did not interfere both the Posner cueing effect and fMRI blood oxygen level-dependent (BOLD) signals in retinotopically organized areas evoked by the target, indicating a gradient or winner-take-all manner, respectively. We also performed a whole-brain group analysis to examine potential cortical or subcortical area(s) that showed a similar interference as those retinotopically organized areas, as well as interregional correlation and intrinsic connectivity analyses to examine the neural loci of saliency map with and without awareness.

## Results

### Psychophysical experiments

In psychophysical experiments, there were four possible foregrounds with 0°, 25°, 50°, and 90° orientation contrasts between the foreground bars and the background bars. In each texture stimulus, a pair of foregrounds was centered in the lower left and lower right quadrants at 5.83° eccentricity (Figure 1A). There were five possible pairs of foregrounds: 90° + 0°, 90° + 25°, 90° + 50°, 25° + 0°, and 50° + 0°. Low- and high-luminance masks (Figure 1B) rendered the whole texture stimulus visible (Experiment 1) or invisible (Experiment 2) to subjects, respectively, confirmed by two-alternative forced choice (2AFC, Experiment 3, *Figure S1A*). In both Experiments 1 and 2, we used a modified version of the Posner paradigm (Posner et al., 1980) to measure the cueing effect induced by the high salient foreground (the target) in each pair of foregrounds, as shown in *Figure 1C*. Namely, the 90°-foreground served as the target for the 90° + 0°, 90° + 25°, and 90° + 50° texture stimuli, the other low salient foreground (i.e., 25° and 50°) that presented at its contralateral counterpart, served as the distractor. Note that there was no distractor for the 90° + 0° texture stimuli since the 0°-foreground region would always contain background bars. Similarly, the 25°- and 50°-foregrounds were the target for the 25° + 0° and 50° + 0° texture stimuli without the distractor, respectively. In our modified Posner paradigm, a valid cue condition was defined as a match of quadrant between the defined target and the ellipse probe (for example, both the 90°-foreground and probe were presented in the lower left quadrant); an invalid cue condition was defined as a mismatch (Figure 1C). Subjects were asked to press one of two buttons as rapidly and correctly as possible to indicate the orientation of the ellipse probe (45° or 135°). There was no significant difference in the false alarm rate, miss rate, or removal rate (i.e., correct reaction times shorter than 200 ms and beyond three standard deviations from the mean reaction time in each condition were removed) across conditions (all *P* > 0.05, partial eta squared, η_p_^2^ < 0.151, *Figure S2*). The cueing effect for each texture stimulus was quantified as the difference between the reaction time of the probe task performance in the invalid cue condition and that in the valid cue condition.

To examine how the saliency map will be distributed among multiple salient regions and how this distribution interacts with awareness, we focused our analysis on the cueing effect of 90°-foreground in three types of texture stimuli: 90° + 0°, 90° + 25°, and 90° + 50° (Figure 1A). In both Visible and Invisible conditions, we found that the 90°-foreground of all the texture stimuli exhibited a positive cueing effect (Visible condition: 90° + 0°: *t*(24) = 17.090, *P* < 0.001, η_p_^2^ = 4.834; 90° + 25°: *t*(24) = 8.338, *P* < 0.001, η_p_^2^ = 2.358; 90° + 50°: *t*(24) = 5.448, *P* < 0.001, η_p_^2^ = 1.541; Invisible condition: 90° + 0°: *t*(24) = 16.038, *P* < 0.001, η_p_^2^ = 4.536; 90° + 25°: *t*(24) = 6.517, *P* < 0.001, η_p_^2^ = 1.843; 90° + 50°: *t*(24) = 10.622, *P* < 0.001, η_p_^2^ = 3.004), indicating that the attention of the subject was attracted more to the 90°-foreground location, allowing them to perform more proficiently in the valid than the invalid cue condition of the probe task (Figure 1D and 1E). Then, these cueing effects were submitted to a repeated measures ANOVA with awareness (Visible and Invisible) and texture stimulus (90° + 0°, 90° + 25°, and 90° + 50°) as within-subjects factors. The main effect of awareness (*F*(1,24) = 41.734, *P* < 0.001, η_p_^2^ = 0.635), the main effect of texture stimulus (*F*(2,48) = 24.258, *P* < 0.001, η_p_^2^ = 0.503), and the interaction between these two factors (*F*(2,48) = 37.159, *P* < 0.001, η_p_^2^ = 0.608) were all significant. Thus, these data were submitted to a further simple effect analysis. Post hoc paired *t* tests showed that the cueing effect in the Visible condition was significantly greater than that in the Invisible condition for both 90° + 0° (*t*(24) = 12.145, *P* < 0.001, η_p_^2^ = 3.123) and 90° + 25° (*t*(24) = 3.396, *P* = 0.002, η_p_^2^ = 0.896) texture stimuli, but not for the 90° + 50° texture stimulus (*t*(24) = −1.334, *P* = 0.195, η_p_^2^ = 0.369). For the Invisible condition, the main effect of the texture stimulus was not significant (*F*(2,48) = 1.252, *P* = 0.287, η_p_^2^ = 0.050, *Figure 1E*), indicating that the cueing effect of 90°-foreground could not be interfered by the distractor (i.e., 25°- and 50°-foregrounds, *Figure 1A*). In other words, the bottom-up attention was completely allocated to the 90°-foreground location (i.e., the winner-take all manner). However, this null interference here could be explained by the distractor that cannot automatically attract subjects’ attention to its location since its low saliency. Accordingly, to address this issue, we analyzed the cueing effect of the 25°- and 50°-foreground within the 25° + 0° and 50° + 0° texture stimuli, respectively. Results against this explanation by showing that the cueing effect of 25°- and 50°-foregrounds were both significantly above zero (25°-foreground: *t*(24) = 3.003, *P* = 0.006, η_p_^2^ = 0.849; 50°-foreground: *t*(24) = 4.185, *P* < 0.001, η_p_^2^ = 1.184, *Figure S3A*).

For the Visible condition, the main effect of the texture stimulus was significant (*F*(2,24) = 47.262, *P* < 0.001, η_p_^2^ = 0.663); post hoc paired *t* tests revealed that the cueing effect of 90° + 25° texture stimulus was significantly lower than that of the 90° + 0° texture stimulus (*t*(24) = 6.113, *P* < 0.001, η_p_^2^ = 1.240) but significantly higher than that of the 90° + 50° texture stimulus (*t*(24) = 3.268, *P* = 0.01, η_p_^2^ = 0.903). These results demonstrated that the cueing effect of 90°-foreground can be interfered by the distractor and, notably, the degree of this interference increased with the orientation contrast of the distractor (i.e., the gradient manner, *Figure 1D*).

In addition, it could be argued that the winner-take-all and gradient manners of saliency map could depend on the strength of cueing effect. To examine this issue, in both Visible and Invisible conditions, the subjects who scored in the top and bottom 48% (i.e., *n* = 12) of the sample’s cueing effect distribution were assigned to High- and Low-cueing effect groups, respectively. Their cueing effects were submitted to a mixed ANOVA with group (High and Low) as the between-subjects factor and texture stimulus (90° + 0°, 90° + 25°, and 90° + 50°) as the within-subjects factor. In both Visible and Invisible conditions, the results showed the same qualitative conclusion and insignificant interactions between the two factors (Figure S4), further confirming that, despite the strength of cueing effect, the bottom-up saliency map was distributed as gradient or winner-take-all manner with or without awareness, respectively.

### fMRI Experiments

Using a block design, the functional magnetic resonance imaging (fMRI) experiment consisted of ten functional runs. Each run consisted of 14 stimulus blocks of 10 s, interleaved with 14 blank intervals of 12 s. There were 14 different stimulus blocks, including 12 different texture stimulus blocks: 3 (texture stimulus: 90° + 0°, 90° + 25°, and 90° + 50°) × 2 (visual field: left and right) × 2 (awareness: Visible and Invisible), and 2 mask-only blocks: low- and high-luminance masks (Figure 2C). Each stimulus block was randomly presented once in each run, and consisted of the same 5 trials. On each trial in the texture stimulus and mask-only blocks, a texture stimulus and the fixation were presented for 50 ms, respectively, followed by a 100-ms mask (low- and high-luminance for Visible and Invisible conditions, respectively) and 1,850-ms fixation. In the Invisible condition, on each trial during both the texture stimulus and mask-only blocks, subjects were asked to press one of two buttons to indicate the location of the 90°-foreground, which was left of fixation in one half of blocks and right of fixation in the other half at random (i.e., the 2AFC task). In the Visible condition, on each trial during the figure block, subjects needed to perform the same 2AFC task of the figure; whereas during the mask-only block, subjects were asked to press one of two buttons randomly (Figure 2C). Behavioral data showed that, our awareness manipulation was effective for both Visible and Invisible conditions (Figure S1C).

**Figure 2.**
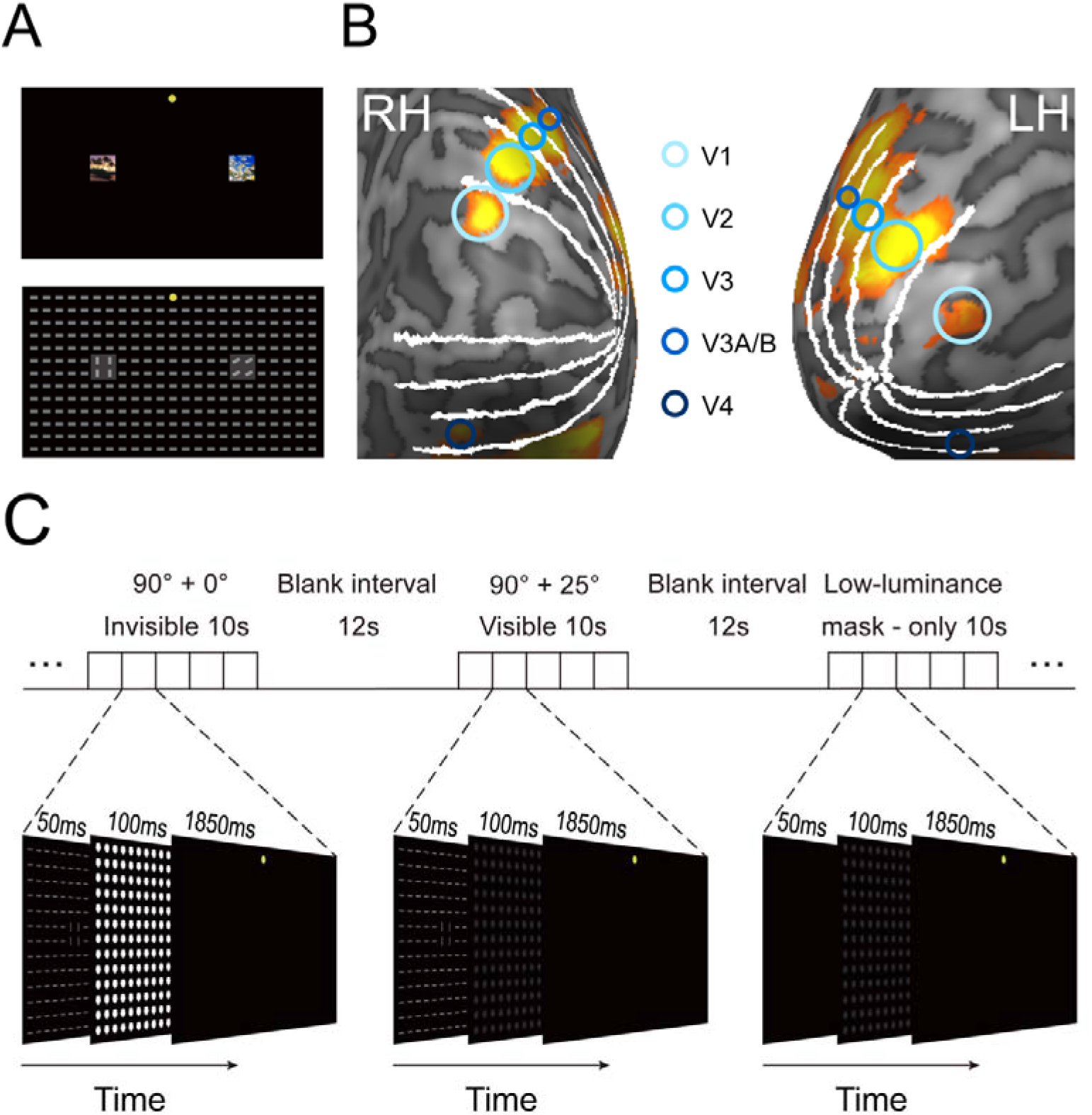
fMRI Stimuli and Protocol. (**A**) ROI definition. The natural scenes in the top panel were used to define ROIs corresponding to the pair of foregrounds in texture stimuli. The transparent squares show the size and location of the natural scenes relative to texture stimuli. (**B**) ROIs on an inflated cortical surface of a representative subject. The ROIs in V1–V4 were defined as the cortical regions responding to the foreground. The boundaries among V1–V4, defined by retinotopic mapping, are indicated by the white lines. (**C**) Block design fMRI procedure. On each trial in the texture stimulus and mask-only blocks, a texture stimulus stimulus and the fixation were presented for 50 ms, respectively, followed by a 100-ms mask (low- and high-luminance for Visible and Invisible conditions, respectively) and 1,850-ms fixation interval. In the Invisible condition, on each trial during both the texture stimulus and mask-only blocks, subjects were asked to press one of two buttons to indicate the location of the 90°-foreground, which was left of fixation in one half of blocks and right of fixation in the other half at random (i.e., the 2AFC task). In the Visible condition, on each trial during the figure block, subjects needed to perform the same 2AFC task of the 90°-foreground; whereas during the mask-only block, subjects were asked to press one of two buttons randomly.

### Region of interest analysis

Regions of interest (ROIs) in SC and V1–V4 were defined as the cortical regions responding significantly to each pair of foregrounds (Figure 2B). BOLD signals were extracted from these ROIs and then averaged according to the stimulus (90° + 0°, 90° + 25°, and 90° + 50° texture stimuli, and the mask-only) and awareness (Visible and Invisible). For each stimulus block, the 2-s preceding the block served as a baseline, and the mean BOLD signal from 5-s to 10-s after stimulus onset was used as a measure of the response amplitude. To isolate the texture stimulus signal, the BOLD amplitudes of the low- and high-luminance mask-only blocks were subtracted from those of the Visible and Invisible texture stimulus blocks, respectively. The BOLD signal difference of the 90°-foreground for each condition is shown in *Figure 3*, and all of them were significantly above zero (all *t*(19) > 2.130, *P* < 0.046, η_p_^2^ > 0.674). Sequentially, in both Visible and Invisible condition, these BOLD signal differences of the 90°-foreground were submitted to a repeated-measures ANOVA with texture stimulus (90° + 0°, 90° + 25°, and 90° + 50°) and cortical area (SC and V1–V4) as within-subjects factors.

**Figure 3.**
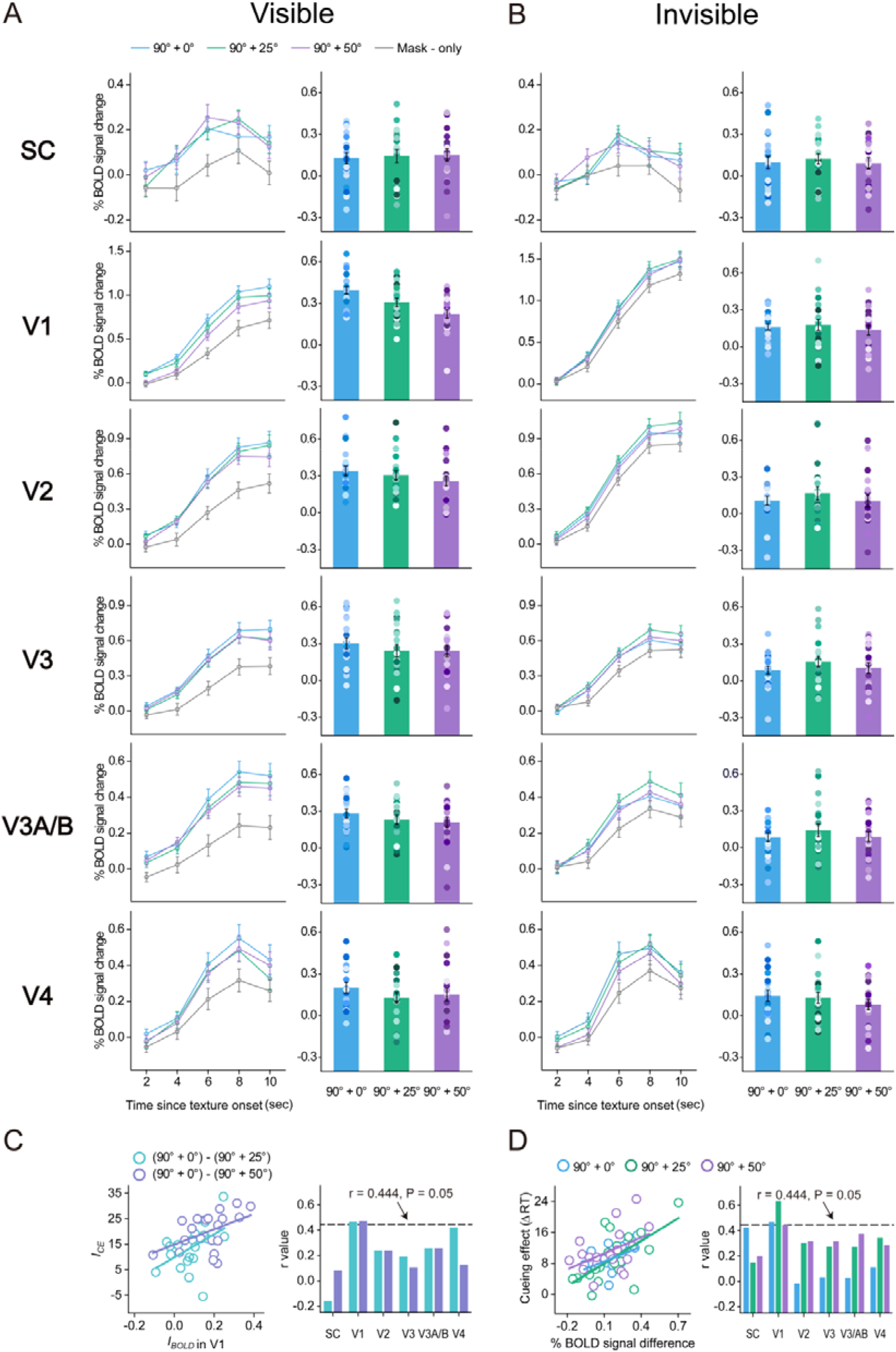
fMRI Results. Left: Blocked BOLD signals averaged across subjects of the ROIs in SC and V1–V4 evoked by the 90°-foreground for three texture stimuli (90° + 0°, 90° + 25°, and 90° + 50°) and the mask-only, during the Visible (**A**) and Invisible (**B**) conditions. Error bars denote 1 SEM calculated across subjects at each time point. Right: BOLD signal differences (i.e., the BOLD signals in the texture stimulus block - the BOLD signals in the mask-only block) of the 90°-foreground in SC and V1–V4 during the Visible (**A**) and Invisible (**B**) conditions. Error bars denote 1 SEM calculated across subjects and colored dots denote the data from each subject. (**C**) Correlations between the *I_CE_* and the *I_BOLD_* in V1 (left), and correlation coefficients (r values) between the *I_CE_* and the *I_BOLD_* in SC and V1–V4 (right), across individual subjects during the Visible condition. (**D**) Correlations between the cueing effect and the BOLD signal difference of 90°-foreground in V1 (left), and correlation coefficients (r values) between the cueing effect and the BOLD signal difference of 90°-foreground in SC and V1–V4 (right), across individual subjects during the Invisible condition.

In the Invisible condition, the main effect of the texture stimulus (*F*(2,38) = 1.100, *P* = 0.342, η_p_^2^ = 0.055), the main effect of cortical area (*F*(5,95) = 0.528, *P* = 0.689, η_p_^2^ = 0.027), and the interaction between these two factors (*F*(10,190) = 0.709, *P* = 0.641, η_p_^2^ = 0.036) were not significant (Figure 3B). These results indicated that the BOLD signal difference of 90°-foreground could not be interfered by the distractor (i.e., 25°- and 50°-foregrounds), i.e., the bottom-up attention was completely allocated to the 90°-foreground location (the winner-take-all manner). Similar to the psychophysical cueing effect (Figure 1E), this null interference could also be explained by the distractor that cannot automatically attract subjects attention to its location since its low saliency. Accordingly, to address this issue, we analyzed the BOLD signal difference of the ROIs evoked by the 25°- and 50°-foregrounds. Results against this explanation by showing that the BOLD signal difference of 25°- and 50°-foregrounds were both significantly above zero (25°-foreground, SC and V1–V4: all *t*(19) > 2.194, *P* < 0.041, η_p_^2^ = 0.694; 50°-foreground, SC and V1–V4: all: *t*(19) > 2.188, *P* < 0.041, η_p_^2^ = 0.692, *Figure S3B*, right). In addition, to examine which cortical area’s activities closely mirrored the psychophysical cueing effect, we calculated the correlation coefficients between the cueing effect and BOLD signal difference of 90°-foreground. The cueing effect was significantly correlated with the BOLD signal difference of the 90°-foreground in V1 (90° + 0°: r = 0.467, *P* = 0.038, η_p_^2^ = 0.218; 90° + 25°: r = 0.633, *P* = 0.003, η_p_^2^ = 0.401; 90° + 50°: r = 0.445, *P* = 0.049, η_p_^2^ = 0.198, *Figure 3D*, left), but not in SC or V2–V4 (90° + 0°: all r < 0.426, *P* > 0.061, η_p_^2^ < 0.181; 90° + 25°: all r < 0.343, *P* > 0.139, η_p_^2^ < 0.118; 90° + 50°: all r < 0.409, *P* > 0.073, η_p_^2^ < 0.167, *Figure 3D*, right). Furthermore, across the three texture stimuli, their mean cueing effect was also significantly correlated with their mean BOLD signal difference of the 90°-foreground in V1 (r = 0.640, *P* = 0.002, η_p_^2^ = 0.410, *Figure S5A*), but not in SC or V2–V4 (all r < 0.315, *P* > 0.176, η_p_^2^ < 0.099, *Figure S5B*). More importantly, the correlation coefficient in V1 was (marginally) significantly larger than those in SC and V2–V4 (*P* = 0.040, 0.060, 0.072, 0.056, and 0.023 for SC, V2, V3, V3A/B, and V4, respectively).

In the Visible condition, the main effect of the texture stimulus was significant (*F*(2,38) = 5.859, *P* = 0.008, η_p_^2^ = 0.236), demonstrating that the BOLD signal difference of 90°-foreground decreased with the orientation contrast of the distractor, namely, increasing the interference of the distractor (the gradient manner). We also found a significant main effect of cortical area (*F*(5,95) = 6.366, *P* < 0.001, η_p_^2^ = 0.251) and a significant interaction between texture stimulus and cortical area (*F*(10, 190) = 2.357, *P* = 0.047, η_p_^2^ = 0.110). Hence, the interference of the distractor decreased gradually from lower to higher cortical areas. This was confirmed in further analysis which showed that the main effect of texture stimulus was significant in V1 (*F*(2,38) = 24.904, *P* < 0.001, η_p_^2^ = 0.567), but not in SC or V2–V4 (all *F*(2,38) < 2.658, *P* > 0.086, η_p_^2^ < 0.123) (Figure 3A). These findings revealed that neural activities in V1 were parallel to the psychophysical cueing effect (Figure 1D). To further evaluate a close relationship between the V1 activities and our psychophysical cueing effect, we computed an interference of the distractor to quantify how much the cueing effect (I_CE_) and BOLD signal (I_BOLD_) of the 90°-foreground changed in both the 90° + 25° and 90° + 50° texture stimuli relative to that in the 90° + 0° texture stimulus. The interference was calculated as follows: *I_CE (90° + 25°)_* = *CE_90° + 0°_* - *CE_90° +_ 25°* and *ICE(90° + 50°)* = *CE90° + 0°* - *CE90° + 50°*, where *CE90° + 0°*, *CE90° +25°*, and *CE90° +50°* are the cueing effect of 90°-foreground for the 90° + 0°, 90° + 25°, and 90° + 50° texture stimuli, respectively. Similarly, for each ROI, *I_BOLD (90° + 25°)_* = *BOLD_90° + 0°_* - *BOLD90° + 25°* and *IBOLD(90° + 50°)* = *BOLD90° + 0°* - *BOLD90° + 50°*, where *BOLD90° + 0°*, *BOLD_90° +25°_*, and *BOLD_90° +50°_* are the BOLD signal difference of 90°-foreground for the 90° + 0°, 90° + 25°, and 90° + 50° texture stimuli, respectively. Subsequently, for each ROI, we calculated the correlation coefficients between the *I_CE_* and the *I_BOLD_* across individual subjects. The results showed that, the *I_CE (90° + 25°)_* and *I_CE (90° + 50°)_* correlated significantly with the *I_BOLD (90° + 25°)_* and *I_BOLD (90° + 50°)_* in V1, respectively (90° + 25°: r = 0.465, *P* = 0.039, η_p_^2^ = 0.216; 90° + 50°: r = 0.472, *P* = 0.036, η_p_^2^ = 0.223, *Figure 3C*, left), but not in other cortical areas (*Figure 3C*, right). Taken together, these results further indicate a close relationship between the V1 activities and psychophysical cueing effect in both visible and invisible conditions.

### Whole-brain group analysis

In both Visible and Invisible conditions, to examine potential cortical or subcortical area(s) whose activities could be modulated by the three types of texture stimuli (90° + 0°, 90° + 25°, and 90° + 50°), we performed a group analysis and did a whole-brain search with a general linear model (GLM) procedure (Friston et al., 1994) for cortical and subcortical area(s) that showed a significant difference among them, after subtracted their respective mask signals (note that the data from the left and right visual fields were combined). Statistical maps were thresholded at p < 0.05 and corrected by FDR correction (Genovese et al., 2002). The results showed that, in the Visible condition, the pIPS (*F*(2,38) = 6.787, *P* = 0.006, η_p_^2^ = 0.263), aIPS (*F*(2,38) = 16.969, *P* < 0.001, η_p_^2^ = 0.472), and FEF (*F*(2,38) = 8.059, *P* = 0.002, η_p_^2^ = 0.298) demonstrated a significant difference among the three types of texture stimuli (Figure 4A). Post hoc paired *t* tests revealed that, for the aIPS, the BOLD signal difference of 90° + 25° texture stimulus was significantly higher than that of the 90° + 0° texture stimulus (*t*(19) = 2.707, *P* = 0.042, η_p_^2^ = 0.210) but significantly lower than that of the 90° + 50° texture stimulus (*t*(19) = −3.500, *P* = 0.007, η_p_^2^ = 0.305); for both the pIPS and FEF, there was no significant difference in the BOLD signal difference between 90° + 25° and 90° + 50° texture stimuli (pIPS: *t*(19) = −0.666, *P* = 1.000, η_p_^2^ = 0.059; FEF: *t*(19) = −1.267, *P* = 0.661, η_p_^2^ = 0.176), and both were significantly higher than 90° + 0° texture stimulus (90° + 25°: pIPS: *t*(19) = 3.409, *P* = 0.009, η_p_^2^ = 0.328; FEF: *t*(19) = 3.090, *P* = 0.018, η_p_^2^ = 0.310; 90° + 50°: pIPS: *t*(19) = 2.859, *P* = 0.030, η_p_^2^ = 0.365; FEF: *t*(19) = 3.910, *P* = 0.003, η_p_^2^ = 0.472). Furthermore, for both 90° + 25° and 90° + 50° texture stimuli, we calculated the correlation coefficients between their *I_BOLD_* in V1 and those in pIPS, aIPS, and FEF across individual subjects. We found that both the *I_BOLD (90° + 25°)_* and *I_BOLD (90° + 50°)_* in V1 correlated significantly with those in pIPS (90° + 25°: r = −0.500, *P* = 0.025, η_p_^2^ = 0.25; 90° + 50°: r = −0.447, *P* = 0.048, η_p_^2^ = 0.20), but not in either aIPS or FEF (Figure 4B). These results indicate a potential involvement of pIPS in the gradient manner of saliency map in V1. In the Invisible condition, however, no such cortical or subcortical areas were found, supporting the idea that the winner-take-all manner of saliency map was constructed in V1.

**Figure 4.**
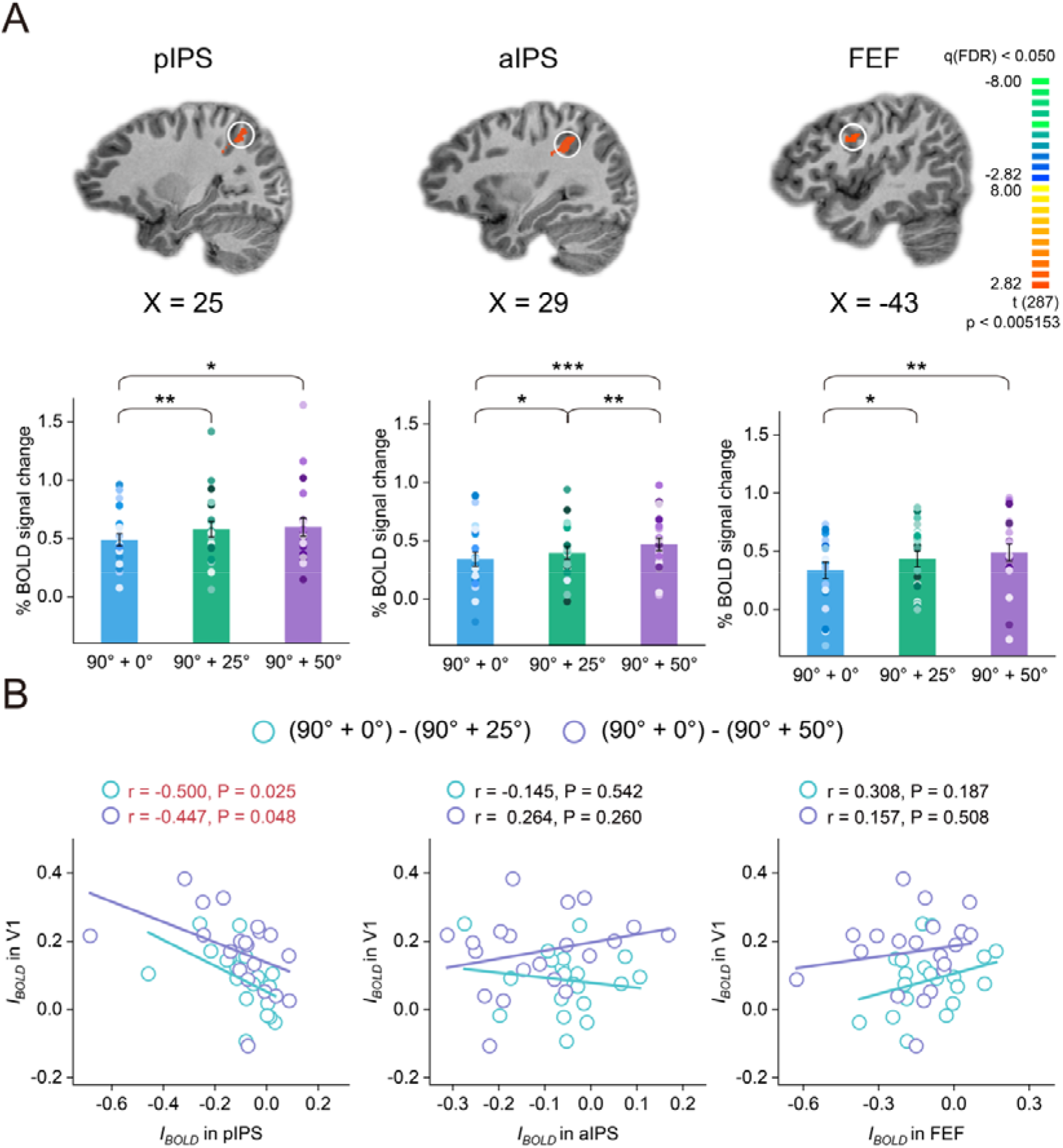
Results of Whole-brain Group Analysis. (**A**) Whole-brain search for pIPS, aIPS, and FEF, with all showing a significant difference in the BOLD signal change among the three types of texture stimuli (90° + 0°, 90° + 25°, and 90° + 50°) during the Visible condition. Note that no such cortical or subcortical areas were found in the Invisible condition (\**P* < 0.05; \**P* < 0.01; \*\*\**P* < 0.001). Error bars denote 1 SEM calculated across subjects and colored dots denote the data from each subject. (**B**) Correlations between the *I_BOLD_* in V1 and that in pIPS (left), aIPS (middle), and FEF (right) across individual subjects during the Visible condition.

### Effective connectivity analysis

In the Visible condition, our results showed that the BOLD signal in both V1 and frontoparietal cortical areas could be modulated by the three types of texture stimuli (90° + 0°, 90° + 25°, and 90° + 50°), indicating a gradient manner of saliency map. To further examine which area is a potential source of this gradient manner of saliency map, we applied dynamic causal modeling (DCM) analysis (Friston, et al., 2003) in SPM12 to examine interregional intrinsic connectivity change among these three types of texture stimuli. Given the extrinsic visual input into both V1-target (i.e., the ROI in V1 was evoked by the target: 90°-foreground) and V1-distractor (i.e., the ROI in V1 was evoked by the distractor), bidirectional intrinsic connections were hypothesized to exist among the pIPS, aIPS, FEF, V1-target, and V1-distractor (*Figure 5A*, top), and these intrinsic connections could be modulated by the texture stimuli.

**Figure 5.**
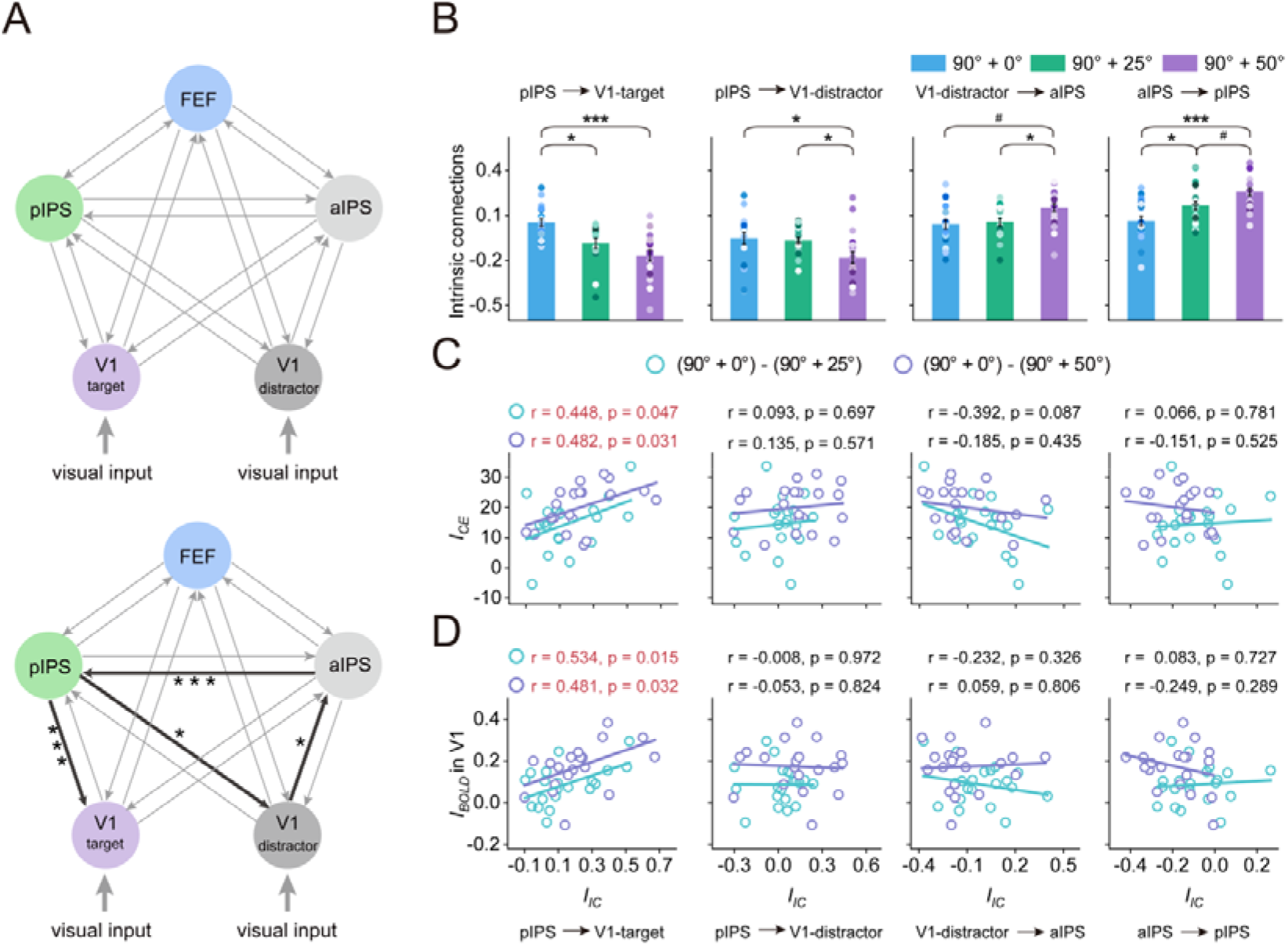
Results of DCM Analysis. (**A**) Top: given the extrinsic visual input into both V1-target (i.e., the ROI in V1 was evoked by the target: 90°-foreground) and V1-distractor (i.e., the ROI in V1 was evoked by the distractor), bidirectional intrinsic connections were hypothesized to exist among the pIPS, aIPS, FEF, V1-target, and V1-distractor. Bottom: thick lines indicate intrinsic connections that are significantly modulated by the three types of texture stimuli (90° + 0°, 90° + 25°, and 90° + 50°) and their significance levels during the Visible condition (\**P* < 0.05; \*\*\**P* < 0.001). (**B**) Intrinsic connections among pIPS, aIPS, V1-target, and V1-distractor for the three types of texture stimuli (^#^0.05 < *P* < 0.07). Error bars denote 1 SEM calculated across subjects and colored dots denote the data from each subject. (**C**) Correlations between the *I_CE_* and the *I_IC_* of each significant intrinsic connection modulated by the three texture stimuli across individual subjects. (**D**) Correlations between the *I_BOLD_* in V1 and the *I_IC_* of each significant intrinsic connection modulated by the three texture stimuli across individual subjects.

Among the 90° + 0°, 90° + 25°, and 90° + 50° texture stimuli, the results showed that, the feedforward connections were significant from V1-distrator (rather than V1-target) to aIPS (*F*(2,38) = 4.327, *P* = 0.023, η_p_^2^ = 0.185), but not to either pIPS or FEF (Figure 5B). Post hoc paired *t* tests revealed that, there was no significant difference between the 90° + 0° and 90° + 25° (*t*(19) = −0.365, *P* = 1.000, η_p_^2^ = 0.121) texture stimuli, and both were (marginally) significantly lower than the 90° + 50° texture stimulus (90° + 0° versus 90° + 50°: *t*(19) = −2.476, *P* = 0.069, η_p_^2^ = −0.859; 90° + 25° versus 90° + 50°: *t*(19) = −2.681, *P* = 0.044, η_p_^2^ = 0.831). These results indicated that the feedforward connections from V1-distractor (rather than V1-target) to aIPS increased with the saliency of distractor. However, the backward connection showed opposite patterns: it was decreased mainly from pIPS to both V1-target (*F*(2,38) = 14.540, *P* < 0.001, η_p_^2^ = 0.434) and V1-distractor (*F*(2,38) = 5.823, *P* = 0.010, η_p_^2^ = 0.434), but not from either aIPS or FEF (Figure 5B). Post hoc paired *t* tests revealed that, for V1-target, there was no significant difference between the 90° + 25° and 90° + 50° (*t*(19) = 2.108, *P* = 0.146, η_p_^2^ = 0.520) texture stimuli, and both were significantly lower than the 90° + 0° texture stimulus (90° + 25° versus 90° + 0°: *t*(19) = −3.317, *P* = 0.011, η_p_^2^ = 1.08; 90° + 50° versus 90° + 0°: *t*(19) = −5.118, *P* < 0.001, η_p_^2^ = 1.534); for V1-distractor, there was no significant difference between the 90° + 0° and 90° + 25° (*t*(19) = 0.455, *P* = 1.000, η_p_^2^ = 0.109) texture stimuli, and both were significantly higher than the 90° + 50° texture stimulus (90° + 0° versus 90° + 50°: *t*(19) = 2.646, *P* = 0.048, η_p_^2^ = 0.758; 90° + 25° versus 90° + 50°: *t*(19) = 2.749, *P* = 0.038, η_p_^2^ = 0.851). In addition, we also found a significant backward connection from aIPS to pIPS (*F*(2,38) = 16.371, *P* < 0.001, η_p_^2^ = 0.463), and post hoc paired *t* tests revealed that the backward connection for the 90° + 25° texture stimuli was (marginally) significantly lower than that for the 90° + 50° texture stimuli (*t*(19) = −2.613, *P* = 0.051, η_p_^2^ = 0.844) but significantly higher than that for the 90° + 0° texture stimuli (*t*(19) = 2.790, *P* = 0.035, η_p_^2^ = 0.689).

To further evaluate the role of these forward and backward connections in the distribution of saliency map, we computed an interference of the distractor to quantify how much the intrinsic connections (I_IC_) changed in the 90° + 0° texture stimulus relative to that in both the 90° + 25° and 90° + 50° texture stimuli. The interference was calculated as follows: *IIC (90° + 25°)* = *IC90° + 0°* - *IC90° + 25°* and *IIC(90° + 50°)* = *IC90° + _0°_* - *IC_90° + 50°_*, where *IC_90° + 0°_*, *IC_90° +25°_*, and *IC_90° +50°_* are the interregional intrinsic connections for the 90° + 0°, 90° + 25°, and 90° + 50° texture stimuli, respectively. Subsequently, for each interregional intrinsic connection and each interference, we calculated the correlation coefficients between the *I_IC_* and the *I_CE_*, and between the *I_IC_* and the *I_BOLD_* across individual subjects. Results showed that, only for the feedback connectivity from pIPS to V1-target, its *I_IC (90° + 25°)_* and *I_IC (90° + 50°)_* correlated significantly with the *I_CE (90° + 25°)_* and *I_CE (90° + 50°)_* in psychophysical experiments (90° + 25°: r = 0.448, *P* = 0.047, η_p_^2^ = 0.201; 90° + 50°: r = 0.482, *P* = 0.031, η_p_^2^ = 0.232, *Figure 5C*), as well as with the *I_BOLD (90° + 25°)_* and *I_BOLD (90° + 50°)_* in V1 (90° + 25°: r = 0.534, *P* = 0.015, η_p_^2^ = 0.285; 90° + 50°: r = 0.481, *P* = 0.032, η_p_^2^ = 0.231, *Figure 5D*), respectively. These results support the idea that the gradient manner of saliency map in V1 was derived by feedback from pIPS rather than from aIPS or FEF.

## Discussion

We examined how the distribution of saliency map interacts with awareness when multiple salient stimuli are presented simultaneously and found the following psychophysical and neuroimaging results. First, we found support for previous neurophysiological (Kastner et al., 1997; Nothdurft et al., 1999; White et al., 2017a; 2017b; Yan et al., 2018), psychophysical (Koene and Zhaoping, 2007; Zhaoping, 2008; Zhaoping and May, 2007; Zhaoping and Zhe, 2015), and brain imaging (Chen et al., 2016; Zhang et al., 2012) studies, indicating that both visible and invisible salient foregrounds could attract bottom-up attention in behavior and evoked greater BOLD signals relative to the background in the texture stimuli. Second, however, there was a critical distinction between the dependence on awareness of them, with visible and invisible salient foregrounds guided bottom-up attention as a gradient or winner-take-all manner, respectively. Finally, the most parsimonious account of our results is that the gradient manner of saliency map in V1 was derived by feedback from pIPS, whereas the winner-take-all manner of saliency map was created in V1.

### Gradient manner of the saliency map in pIPS

In accordance with previous studies demonstrating a direct representation for pIPS of the saliency map (Bisley and Goldberg, 2010; Bogler et al., 2011; Buschman and Miller, 2007; Constantinidis and Steinmetz, 205; Gottlieb, 2007; Gottlieb et al., 1998; Serences et al., 2005), our study further revealed that pIPS could control the gradient manner of saliency map in V1. One should note that the gradient manner in our study was reflected by the decreased response to the 90°-foreground with the increased orientation contrast of the distractor. Thus, how does pIPS decrease V1 responses to the salient foreground? Previous studies have implicated IPS in the suppression of task-irrelevant distractors (Mevorach et al., 2009, 2010; Payne and Allen, 2011; Ruff and Driver, 2006), and our findings are consistent with such an influence. First, pIPS responses were significantly modulated by the texture stimuli (90° + 0°, 90° + 25°, and 90° + 50°, *Figure 4A*) and showed a gradient pattern of activation consistent with that in V1 (Figure 3A), where the BOLD signal of 90°-foreground decreased (i.e., the increased interference) with the orientation contrast of the distractor. Second, these increased interferences in V1 were significantly predicted by the enhanced response in pIPS rather than in other frontoparietal cortical areas (Figure 4B). Finally, the DCM analysis indicated that the distractor significantly modulated the feedback from pIPS to V1 (rather than the feedforward from V1 to pIPS, *Figure 5A*, bottom), and the decreased feedback significantly predicted the increased interference of distractor in V1 (Figure 5D) and that in psychophysical cueing effect (Figure 5C).

Our findings can be viewed as identifying the human parietal cortex as a source of the gradient manner of saliency map. Note that, this conclusion is based mainly on our DCM analyses, which depended on time-series models of fMRI data for an interpretation of causality (Friston et al., 2003). The interpretation of causality in our study finds support in previous lesion (Corbetta and Shulman, 2011; Shomstein, 2012) and transcranial magnetic stimulation (Hodsoll et al., 2009; Mevorach et al., 2006) studies showing a causal effect of parietal cortical disruption on bottom-up attention driven by the salient stimulus. The prominent role of the parietal cortex in the gradient manner of saliency map evident here not only is consistent with recent neurophysiological findings that have revealed how parietal areas directly realize the saliency map (Bisley and Goldberg, 2010; Buschman and Miller, 2007; Constantinidis and Steinmetz, 2005; Gottlieb, 2007; Gottlieb et al., 1998), but also support the dominant model of the saliency map developed by Itti and Koch (2001), which proposes that higher cortical areas, particularly the parietal and frontal cortex, whose neurons are less selective to specific visual features (i.e., color, orientation, or other features, Koch and Ullman, 1985; Wolfe, 1994), are more likely to be possible candidates that realize the saliency map.

Although we emphasize the importance of pIPS in the gradient manner of saliency map, we cannot deny a potential contribution from other parietal cortex, such as aIPS. Actually, previous neurophysiological and brain imaging studies (Bogler et al., 2011; Geng and Mangun, 2009; Serences and Yantis, 2007) have reported that the aIPS could also represent the saliency map. Our study supported these findings by showing that the BOLD signal in aIPS (Figure 4A) and the feedforward from V1 (in particular, V1-distractor) to aIPS (Figure 5B) were significantly modulated by the texture stimuli. However, more importantly, our results showed that the feedback from aIPS to V1 (including both the V1-target and V1-distractor) was not modulated by the texture stimuli (Figure 5B), indicating that the aIPS is more likely to inherit or read out saliency signals from V1 or other earlier areas. Interestingly, we also found a significant backward connection from aIPS to pIPS in our study (Figure 5B). This backward connection, however, could not significantly predict either the BOLD signal change in V1 (Figure 5D) or the cueing effect in psychophysical experiments (Figure 5C). These results suggest that the involvement of aIPS in the gradient manner of saliency map in our study may occur via pIPS. Indeed, using multivariate pattern analysis of fMRI, previous studies have indicated that the pIPS and aIPS may be involved in different stages of saliency map (Bogler et al., 2011; Itti and Koch, 2001). Given the low temporal resolution of fMRI, further work is needed using neurophysiological techniques to parse how these two areas are involved in the distribution of saliency map.

### Winner-take-all manner of the saliency map in V1

Compared to a gradient manner of saliency map with awareness in pIPS, our results suggest that the saliency map without awareness is distributed as a winner-take-all manner and this manner is created in V1. Our claim was based on the following findings. First, both the cueing effects (Figure 1E) and BOLD signal changes (Figure 3B) of 90°-foreground were not interfered by the distractor (25°- and 50°-foregrounds) as showing a null difference among 90° + 0°, 90° + 25°, and 90° + 50° texture stimuli. In other word, the bottom-up attention, despite of the distractor, was completely allocated to the 90°-foreground location. Second, the psychophysical cueing effect was significantly correlated with the BOLD signal change in V1 (*Figure 3D*, left), but not elsewhere (*Figure 3D*, right). More importantly, such correlation coefficient in V1 was (marginally) significantly larger than those in other cortical areas (Figure S5). Finally, using a voxel-wise analysis, we didn’t find any cortical or subcortical areas that modulated by the distractor, demonstrating that the observed neural activities in V1 were not attributed to signals either feedforward or feedback from other areas. Notably, we acknowledge that this winner-take-all manner of saliency map relies on a null difference among the three texture stimuli. However, the 90°-foreground in all the texture stimuli showed significant cueing effects (Figure 1E) and BOLD signal changes (Figure 3B). Hence, our results reflect positive findings. Moreover, it could be argued that this winner-take-all manner can also be explained by the distractor (25°- and 50°-foregrounds) that cannot automatically attract subjects attention to its location since its low saliency. Our results against this explanation by showing that both the cueing effect and BOLD signal changes of distractors were significantly above zero (*Figure S3B*, right).

Our study succeeded in linking V1 activities directly with the winner-take-all manner of saliency map. These findings not only are consistent with existing psychophysical (Koene and Zhaoping, 2007; Zhaoping, 2008; Zhaoping and May, 2007; Zhaoping and Zhe, 2015), neurophysiological (Kastner et al., 1997; Nothdurft et al., 1999; Yan et al., 2018), and brain imaging (Chen et al., 2016; Zhang et al., 2012) studies suggesting that V1 realizes a purely saliency map, but also support and extend the V1 saliency theory (Li, 1999, 2002). This theory proposes that, saliency of a visual location is determined by its highest evoked V1 response relative to those evoked by other locations. In other words, saliency is determined by the relative rather than absolute levels of V1 responses. The dependence of saliency on the relative rather than the absolute levels of neural responses are strongly compatible with our results here. The 90°-foreground within all texture stimuli, despite of the saliency of distractor, always has the highest saliency and thus can completely attract subjects’ attention to its location, i.e., the winner-take-all manner.

### Awareness-dependent distribution of the saliency map

Previous studies have reported ostensibly conflicting results with regard to the neural substrate of saliency map. Three of our previous studies (Chen et al., 2016; Huang et al., 2020; Zhang et al., 2012) suggest that discrepancies in the literature findings may have resulted from the possible contamination by top-down signals, specifically the conscious access to salient stimuli, which have not been systematically controlled or manipulated. Here our study addresses it using a backward masking paradigm in which the low- and high-luminance mask renders the salient foreground (and indeed the whole texture stimuli) visible and invisible to subjects, respectively. We assume that relative to the visible foreground, the invisible foreground can maximally reduce various top-down contaminations, such as feature perception, object recognition, and subjects’ intentions (Zhang et al., 2012). It has been proved and widely accepted that subjective awareness is determined by top-down signaling (Del Cul et al. 2007; Mashour et al., 2020). Thus, rendering a stimulus invisible could maximally (although not completely, Boly et al., 2017) reduce top-down signals that evoked by stimuli, especially for temporally sluggish fMRI signals that typically reflect neural activities resulting from both bottom-up and top-down processes (Fang et al., 2008). We thus speculate that the higher cortical areas, particularly the parietal and frontal cortex, are more likely to be the neural substrate of saliency map with the visible stimulus, since these areas are able to integrate top-down and bottom-up attention (Bisley and Goldberg, 2010; Bogler et al., 2011; Geng and Mangun, 2009; Gottlieb et al., 1998; Katsuki and Constantinidis, 2012; Squire et al., 2013). Conversely, early visual areas are more likely to be possible candidates that realize a purely saliency map of the invisible stimulus, specifically the area V1 that is nearly independent of top-down modulation. Indeed, previous studies have demonstrated that top-down control modulates extrastriate areas but not V1 (Kastner et al. 1998; Luck et al. 1997; Melloni et al., 2012) and decreases gradually from extrastriate visual areas to V1 (Buffalo et al. 2010; Liu et al., 2005). Accordingly, our findings support these speculations and identify the parietal cortex and V1 as the neural substrate of saliency map with and without awareness, respectively.

In addition, at least two intriguing questions need to be addressed in further research. First, it should be noted that the awareness-dependent distribution of saliency map in our study was indexed by the interference of the distractor (25°- and 50°-foregrounds), which was always presented in an opposite visual field to the target (90°-foreground) and presumably induced the divided attention. Interestingly, a number of previous studies have demonstrated that the divided attention is modulated by whether targets and/or distractors are presented across hemifields or within the same hemifield (Cavanagh and Alvarez, 2005; Franconeri et al., 2013). In other words, the interference of distractor (i.e., the awareness-dependent distribution of saliency map) could be modulated by whether the target (90°-foreground) and distractor (25°- and 50°-foregrounds) are presented across or within the same hemifield, and thus further studies will shed light on this issue. Second, compared with current stimuli that consist of simple oriented bars (Figure 1A), complex natural scenes that contain richer naturalistic low-level features (e.g., luminance, contrast, orientation, spatial frequency, and curve, all of which are highly tuned by the visual system), are also thought to be optimal for automatic bottom-up attention (Bogler et al. 2011; Chen et al., 2016; White et al., 2017a). Further work is thus needed to address whether our conclusion can be generalized to the complex natural scene.

### Conclusions

We conclude that, when multiple salient stimuli are presented simultaneously, the distribution of saliency map is either a gradient or winner-take-all manner, depending on the conscious access to salient stimuli. Our study provides, to the best of our knowledge, the first evidence for awareness-dependent saliency map and its distinct neural loci with and without awareness. Identifying the parietal cortex and V1 as the neural substrate of saliency map with and without awareness, respectively, reconciles previous, seemingly contradictory findings on the saliency map.

## Materials and Methods

### Subjects

A total of 25 human subjects (4 male, 19–26 years old) were involved in the study. All of them participated in the psychophysical experiment. Twenty-one of them participated in the fMRI experiment. One subject in the fMRI experiment was excluded because of large head motion (> 3 mm). They were naïve to the purpose of the study. They were right-handed, reported normal or corrected-to-normal vision, and had no known neurological or visual disorders. They gave written, informed consent, and our procedures and protocols were approved by the human subjects review committee of School of Psychology at South China Normal University.

### Stimuli

Each texture stimulus (Figure 1A) had a regular Manhattan grid of 13 × 25 low luminance bars (1.38 cd/m^2^), presented in the lower visual field on a dark screen (0.007 cd/m^2^). Each bar was a rectangle of 0.0625° × 0.5° in visual angle. The center-to-center distance between the bars was 0.75°. All bars were identically oriented except for a foreground region of 2 × 2 bars with another orientation. The orientation of the background bars was randomly chosen from 0° to 180° on each trial. There were four different foregrounds with 0°, 25°, 50°, and 90° orientation contrasts between the foreground bars and the background bars (Figure 1A). In each texture stimulus, a pair of foregrounds was centered in the lower left and lower right quadrants at 5.83° eccentricity. There were five possible pairs of foregrounds: 90° + 0°, 90° + 25°, 90° + 50°, 25° + 0°, and 50° + 0° in our experiments. Low- (0.016 cd/m^2^) and high- (78.675 cd/m^2^) luminance masks, which had the same grid as the texture stimuli, rendered the whole texture stimulus visible (Experiment 1) and invisible (Experiment 2, confirmed by a 2AFC test, i.e., Experiment 3) to subjects, respectively. Each element of the mask contained 12 intersecting bars oriented from 0° to 165° at every 15° interval. The bars in the mask had the same size and shape as those in the texture stimuli.

### Psychophysical experiments

Visual stimuli were displayed on an IIYAMA color graphic monitor (model: HM204DT; refresh rate: 60 Hz; resolution: 1,280 × 1,024; size: 22 inches) at a viewing distance of 57 cm. Subjects’ head position was stabilized using a chin rest. A yellow fixation point was always present at the center of the monitor.

Psychophysical experiments consisted of three experiments. Experiments 1 (Visible) and 2 (Invisible) investigated whether the distribution of saliency map depended on the visibility of texture stimuli. Subjects participated in Experiments 1 and 2 on two different days, and the order of the two experiments was counterbalanced across subjects. In both Experiments 1 and 2, we used a modified version of the Posner paradigm (Posner et al., 1980) to measure the spatial cueing effect induced by the high salient foreground (the target, *Figure 1C*). Namely, the 90°-foreground served as the target for the 90° + 0°, 90° + 25°, and 90° + 50° texture stimuli, the other low salient foreground (i.e., 25° and 50°) that presented at its contralateral counterpart, served as the distractor. Note that there was no distractor for the 90° + 0° texture stimuli since the 0°-foreground region would always contain background bars. Similarly, the 25°- and 50°-foregrounds were the target for the 25° + 0° and 50° + 0° texture stimuli without the distractor, respectively. Each trial began with the fixation. A texture stimulus was presented for 50 ms, followed by a 100-ms mask (low- and high-luminance in Experiments 1 and 2, respectively) and another 50-ms fixation interval. The defined target in the texture stimulus served as a cue to attract spatial attention. Then an ellipse probe was presented for 50-ms at randomly either the target location (e.g., the 90°-foreground location, valid cue condition) or its contralateral counterpart (the distractor location, i.e., invalid cue condition) with equal probability (Figure 1C). The ellipse probe was orientated at 45° or 135° away from the vertical. Subjects were asked to press one of two buttons as rapidly and correctly as possible to indicate the orientation of the ellipse probe (45° or 135°). Each experiment consisted of 12 blocks, 6 for the 25°-distractor and 6 for the 50°-distractor. The order of these two different distractors was counterbalanced across subjects. Each block had 96 trials, from randomly interleaving 32 trials from three conditions (25°-distractor: 90° + 0°, 90° + 25°, and 25° + 0°; 50°-distractor: 90° + 0°, 90° + 50°, and 50° + 0°). The cueing effect for each texture stimulus was quantified as the difference between the reaction time of the probe task performance in the invalid cue condition and that in the valid cue condition.

Experiment 3 checked the effectiveness of the awareness manipulation in Experiments 1 and 2, and was always before them. In Experiment 3, all subjects underwent a 2AFC task to determine whether the masked foreground was visible or invisible in a criterion-free way. The stimuli and procedure in this 2AFC experiment were the same as those in Experiments 1 and 2, except that no probe was presented (Figure S1A). After the presentation of a masked texture stimulus, subjects were asked to make a forced choice response regarding which side (lower left or lower right) from the fixation they thought the defined target appeared. Their performances were significantly higher or not statistically different from chance for all possible texture stimuli, providing an objective confirmation that the foreground was indeed visible or invisible to subjects, respectively.

### fMRI experiments

Using a block design, the experiment consisted of 10 functional runs. Each run consisted of 14 stimulus blocks of 10 s, interleaved with 14 blank intervals of 12 s. There were 14 different stimulus blocks: 3 (texture stimulus: 90° + 0°, 90° + 25°, and 90° + 50°) × 2 (visual field: left/right) × 2 (awareness: visible/invisible), and 2 mask-only blocks: low- and high-luminance masks. Each stimulus block was randomly presented once in each run, and consisted of 5 trials. On each trial in the texture stimulus and mask-only blocks, a texture stimulus or a fixation was presented for 50 ms, respectively, followed by a 100-ms mask (low- and high-luminance for Visible and Invisible conditions, respectively) and 1,850-ms fixation (Figure 2C). In the Invisible condition, on each trial during both the texture stimulus and mask-only blocks, subjects were asked to press one of two buttons to indicate the location of the 90°-foreground (the target), which was left of fixation in one half of blocks and right of fixation in the other half at random (i.e., the 2AFC task). Note that although the salient foreground wasn’t presented in the mask-only blocks, subjects also indicated the location of the target since in the invisible condition; they were unaware whether the target was present or absent. In the Visible condition, on each trial during the texture stimulus block, subjects needed to perform the same 2AFC task of the target; whereas during the mask-only block, subjects were asked to press one of two buttons randomly since in the visible condition, they were easy to perceive the absence of targets.

Retinotopic visual areas (SC and V1–V4) were defined by a standard phase-encoded method developed by Sereno et al. (1995) and Engel et al. (1997), in which subjects viewed rotating wedge and expanding ring stimuli that created traveling waves of neural activity in visual cortex. An independent block-design scan was used to localize the ROIs in SC and V1–V4 corresponding to the pair of foreground regions. The scan consisted of 12 12-s stimulus blocks, interleaved with 12 12-s blank intervals. In a stimulus block, subjects passively viewed images of colorful natural scenes, which had the same size as the foreground regions in texture stimuli and were presented at locations of the pair of foreground regions. Images appeared at a rate of 8 Hz.

### MRI data acquisition

MRI data were collected using a 3T Siemens Trio scanner with a 32-channel phase-array coil at the Center for MRI Research at South China Normal University. In the scanner, the stimuli were back-projected via a video projector (refresh rate: 60 Hz; spatial resolution: 1,024 × 768) onto a translucent screen placed inside the scanner bore. Subjects viewed the stimuli through a mirror located above their eyes. The viewing distance was 90 cm. Blood oxygen level-dependent (BOLD) signals were measured with an echo-planar imaging sequence (TE: 30 ms; TR: 2000 ms; FOV: 196 × 196 mm^2^; matrix: 128 × 128; flip angle: 90; slice thickness: 3 mm; gap: 0 mm; number of slices: 32, slice orientation: axial). A high-resolution 3D structural data set (3D MPRAGE; 1 × 1 × 1 mm^3^ resolution; TR: 2600 ms; TE: 3.02 ms; FOV: 256 × 256 mm^2^; flip angle: 8; number of slices: 176; slice orientation: sagittal) was collected in the same session before the functional scans. Subjects underwent three sessions, one for the retinotopic mapping and ROI localization, and the other two for the main experiment.

### MRI data analysis

Note that, the MRI data analysis, whole-brain group analysis, and DCM of this study closely followed those used by our previous studies (Zhang et al., 2014, 2016, 2018) and therefore, for consistency, we largely reproduce that description here, noting differences as necessary. The anatomical volume for each subject in the retinotopic mapping session was transformed into a brain space that was common for all subjects (Talairach and Tournoux, 1988) and then inflated using BrainVoyager QX. Functional volumes in both sessions for each subject were preprocessed, including 3D motion correction, linear trend removal, and high-pass (0.015 Hz, Smith et al., 1999) filtering using BrainVoyager QX. Head motion within any fMRI session was < 3 mm for all subjects. The images were then aligned to the anatomical volume in the retinotopic mapping session and transformed into Talairach space (Talairach and Tournoux, 1988). The first 8-s of BOLD signals were discarded to minimize transient magnetic saturation effects.

A general linear model (GLM) procedure was used for the ROI analysis. The ROIs in SC and V1–V4 were defined as areas that responded more strongly to the natural scene images than blank screen (*P* < 0.05, corrected by FDR correction, Genovese et al., 2002). The block-design BOLD signals were extracted from these ROIs and then averaged according to each type of trials. For each stimulus block, the 2-s preceding the block served as a baseline, and the mean BOLD signal from 5-s to 10-s after stimulus onset was used as a measure of the response amplitude.

### Whole-brain group analysis

In the whole-brain group analysis, a fixed-effects general linear model (FFX-GLM) was performed for each subject on the spatially non-smoothed functional data in Talairach space. The 1^st^-level regressors were created by convolving the onset of each stimuli block with the default BrainVoyager QX‘s two-gamma hemodynamic response function. Six additional parameters resulting from 3D motion correction (x, y, z rotation and translation) were included in the model. In both Visible and Invisible conditions, for each subject, we first calculated fixed effects analyses for each texture stimulus separately (90° + 0° texture stimulus block vs. mask-only block, 90° + 25° texture stimulus block vs. mask-only block, and 90° + 50° texture stimulus block vs. mask-only block) (Figure 2C). Next, a second-level group analysis (n = 20) was performed with a random-effects GLM to calculate the contrast among the 90° + 0°, 90° + 25°, and 90° + 50° texture stimuli. Statistical maps were thresholded at *P* < 0.05 and corrected by FDR correction (Genovese et al., 2002).

### Effective connectivity analysis

In the Visible condition, our results showed that the BOLD signal in both V1 and frontoparietal cortical areas could be modulated by the three types of texture stimuli (90° + 0°, 90° + 25°, and 90° + 50°), indicating a gradient manner of saliency map. To further examine which area is a potential source of this gradient manner of saliency map, we applied DCM analysis (Friston, et al., 2003) in SPM12 to examine interregional intrinsic connectivity change among these three types of texture stimuli. For each subject and each hemisphere, using BrainVoyager QX, V1 voxels were identified as those activated by the foreground at a significance level of p < 0.01. All of the pIPS, aIPS, and FEF voxels were identified as those activated by the stimulus block at a significance level of p < 0.01. The mean Talairach coordinates of V1, pIPS, aIPS, and FEF, and their standards errors across subjects were [-10 ± 0.99, −95 ± 0.96, 0 ± 1.74], [-25 ± 1.04, −65 ± 1.04, 40 ± 0.98], [-39 ± 1.70, −44 ± 1.87, 38 ± 4.71], and [-46 ± 0.82, 2 ± 0.94, 31 ± 0.90] for the left hemisphere, and [7 ± 0.98, −93 ± 0.63, 3 ± 1.57], [25 ± 0.81, −64 ± 1.46, 40 ± 1.79], [31 ± 1.16, −45 ± 1.06, 39 ± 0.90], and [44 ± 0.57, 3 ± 0.57, 24 ± 1.01], for the right hemisphere, respectively. For each subject and each hemisphere, these Talairach coordinates were converted to Montreal Neurological Institute (MNI) coordinates using the tal2mni conversion utility (http://imaging.mrc-cbu.cam.ac.uk/downloads/MNI2tal/tal2mni.m). In SPM, for each of these areas, we extracted voxels within a 6-mm sphere centered on the most significant voxel and used their time series for the DCM analysis. The estimated DCM parameters were later averaged across the two hemispheres using the Bayesian model averaging method (Penny et al., 2004).

Given the extrinsic visual input into both V1-target (i.e., the ROI in V1 was evoked by the target: 90°-foreground) and V1-distractor (i.e., the ROI in V1 was evoked by the distractor), bidirectional intrinsic connections were hypothesized to exist among the pIPS, aIPS, FEF, V1-target, and V1-distractor (*Figure 5A*, top), and these intrinsic connections could be modulated by the texture stimuli (90° + 0°, 90° + 25°, and 90° + 50°).

## Acknowledgements

We thank Li Zhaoping for valuable comments. This work was supported by the National Natural Science Foundation of China (Projects 32022032 and 31871135) and the Key Realm R&D Program of Guangzhou (202007030005).

## Additional information

### Funding

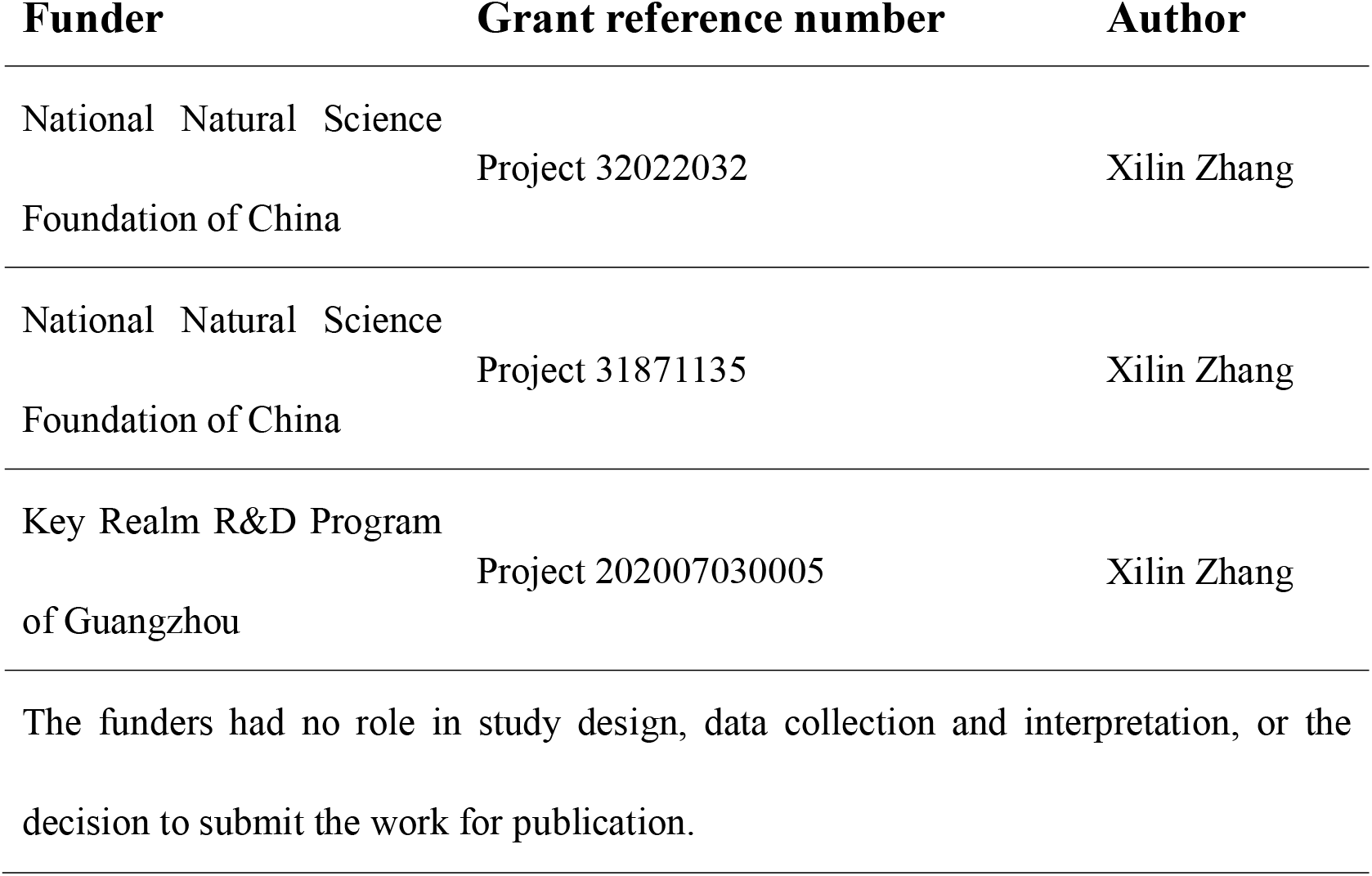

## Author contributions

Lijuan Wang, Ling Huang, Mengsha Li, Xiaotong Wang, Shiyu Wang, Yuefa Lin, Conceptualization, Formal analysis, Investigation, Methodology, Writing – original draft, Writing - review and editing; Xilin Zhang, Conceptualization, Formal analysis, Supervision, Funding acquisition, Investigation, Methodology, Writing - original draft, Writing - review and editing

### Ethics

Human subjects: They study was approved by the human subjects review committee of School of Psychology at South China Normal University.

## Additional files

### Data availability

The code and MRI dataset generated during this study is available on Open Science Framework: https://osf.io/u7sbf/.

## Supplementary Information

**Figure S1.**
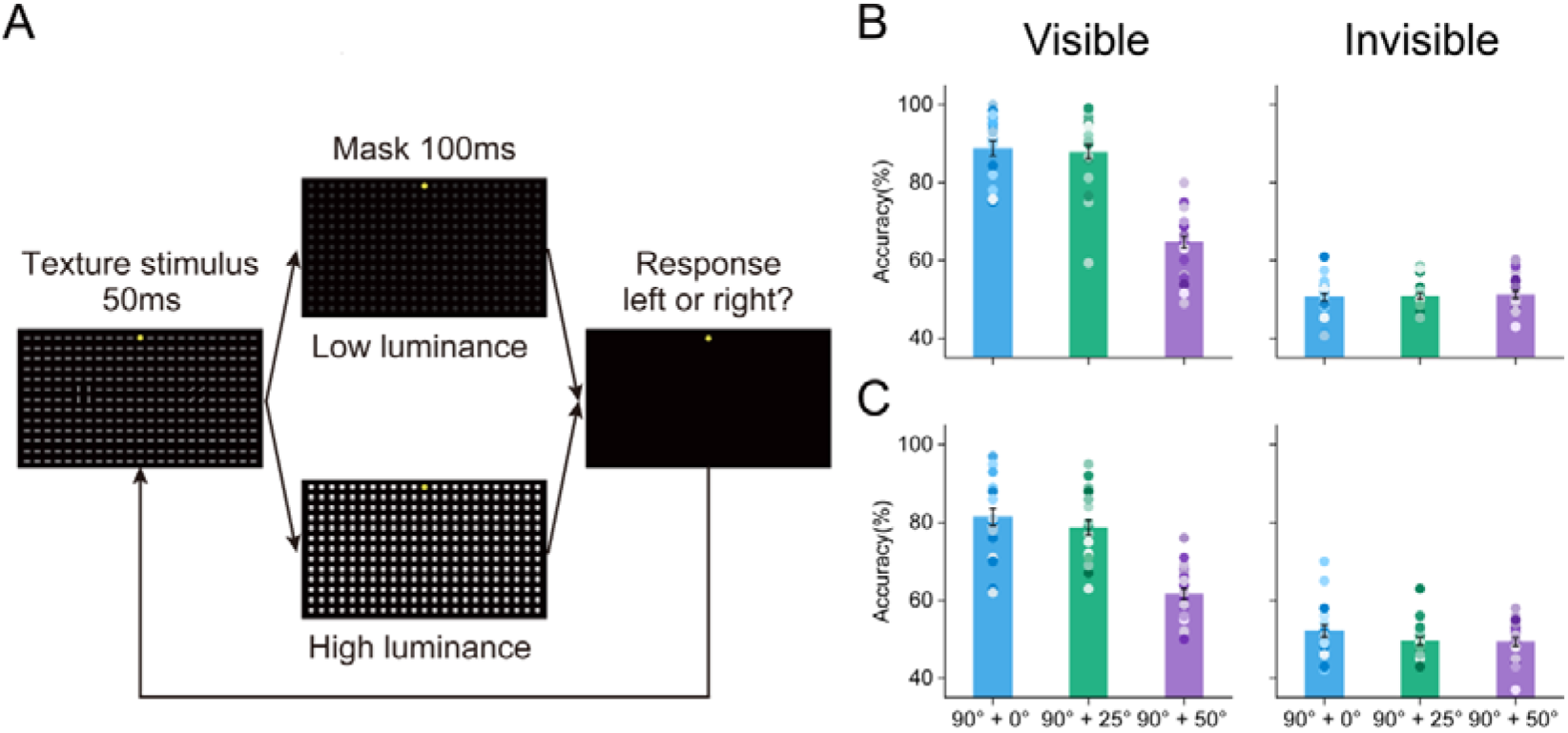
Protocol and Results of the 2AFC Test in Psychophysical Experiments, and Behavioral Data in fMRI Experiments. (**A**) The stimuli and procedure in the 2AFC experiment (i.e., Experiment 3) were the same as those in Experiments 1 and 2, except that no probe was presented. After the presentation of a masked texture stimulus, subjects were asked to make a forced choice response regarding which side (lower left or lower right) from the fixation they thought the 90°-foreground appeared. (**B**) In the psychophysical experiments, for the Invisible condition, subjects reported that they were unaware of the texture stimuli and could not detect which quadrant contained the 90°-foreground in psychophysical experiments. Their performances were not statistically different from chance (mean percent correct ± standard error of the mean [SEM], 90° + 0°: 50.609 ± 0.893%, *t*(24) = 0.683, *P* = 0.501, η_p_^2^ = 0.193; 90° + 25°: 50.906 ± 0.780%, *t*(24) = 1.163, *P* = 0.256, η_p_^2^ = 0.329; 90° + 50°: 51.230 ± 0.926%, *t*(24) = 1.328, *P* = 0.197, η_p_^2^ = 0.376); for the Visible condition, by contrast, their performance was significantly higher than chance (90° + 0°: 88.715 ± 1.920%, *t*(24) = 20.162, *P* < 0.001, η_p_^2^ = 5.703; 90° + 25°: 87.929 ±1.845%, *t*(24) = 20.563, *P* < 0.001, η_p_^2^ = 5.816; 90° + 50°: 64.753 ± 1.562%, *t*(24) = 9.442, *P* < 0.001, η_p_^2^ = 2.671), indicating that our awareness manipulation was effective for both Visible and Invisible conditions. Error bars denote 1 SEM calculated across subjects and colored dots denote the data from each subject. (**C**) In fMRI experiments, for the Invisible condition, on each trial during both the texture stimulus and mask-only blocks, subjects were asked to press one of two buttons to indicate the location of the 90°-foreground, which was left of fixation in one half of blocks and right of fixation in the other half at random (i.e., the 2AFC task). For the Visible condition, on each trial during the texture stimulus block, subjects needed to perform the same 2AFC task of the 90°-foreground; whereas during the mask-only block, subjects were asked to press one of two buttons randomly. Note that in both Visible and Invisible conditions, we only calculated subjects performances during the texture stimulus blocks. Behavioral data showed that, in the Invisible condition, subjects’ performances were not statistically different from chance (90° + 0°: 52.100 ± 1.501%, *t*(19) = 1.399, *P* = 0.178, η_p_^2^ = 0.443; 90° + 25°: 49.550 ± 1.045%, *t*(19) = −0.431, *P* = 0.672, η_p_^2^ = 0.136; 90° + 50°: 49.350 ± 1.152%, *t*(19) = −0.564, *P* = 0.579, η_p_^2^ = 0.178), in the Visible condition, by contrast, their performances were significantly higher than chance (90° + 0°: 81.400 ± 2.200%, *t*(19) = 14.274, *P* < 0.001, η_p_^2^ = 4.514; 90° + 25°: 78.750 ± 1.971%, *t*(19) = 14.589, *P* < 0.001, η_p_^2^ = 4.613; 90° + 50°: 61.650 ± 1.466%, *t*(19) = 7.947, *P* < 0.001, η_p_^2^ = 2.513). Furthermore, for both psychophysical and fMRI experiments, subjects performances were submitted to a repeated-measures ANOVA with awareness (Visible and Invisible) and texture stimulus (90° + 0°, 90° + 25°, and 90° + 50°) as within-subjects factors. For both psychophysical and fMRI experiments, the main effect of awareness (the psychophysical experiment: *F*(1, 24) = 314.083, *P* < 0.001, η_p_^2^ = 0.929; the fMRI experiment: *F*(1, 19) = 247.918, *P* < 0.001, η_p_^2^ = 0.929), the main effect of texture stimulus (the psychophysical experiment: *F*(2, 48) = 73.596, *P* < 0.001, η_p_^2^ = 0.754; the fMRI experiment: *F*(2, 38) = 74.40, *P* < 0.001, η_p_^2^ = 0.797), and the interaction between the two factors (the psychophysical experiment: *F*(2, 48) = 83.479, *P* < 0.001, η_p_^2^ = 0.777; the fMRI experiment: *F*(2, 38) = 41.390, *P* < 0.001, η_p_^2^ = 0.685) were all significant. Thus, these data were submitted to a further simple effect analysis. Post hoc paired *t* tests showed that subjects’ performances in the Visible condition was significantly greater than those in the Invisible condition for all three texture stimuli (the psychophysical experiment: all *t*(24) > 6.595, *P* < 0.001, η_p_^2^ > 2.106; the fMRI experiment: all *t*(24) > 8.540, *P* < 0.001, η_p_^2^ > 2.086). For the Invisible condition, the main effect of the texture stimulus was not significant in either psychophysical (*F*(2, 48) = 0.178, *P* = 0.822, η_p_^2^ = 0.007) or fMRI (*F*(2, 38) = 2.669, *P* = 0.086, η_p_^2^ = 0.123) experiments. For the Visible condition, however, the effect of the texture stimulus was significant in both psychophysical (*F*(2, 48) = 101.715, *P* < 0.001, η_p_^2^ = 0.809) and fMRI (*F*(2, 38) = 95.065, *P* < 0.001, η_p_^2^ = 0.833) experiments; Post hoc paired *t* tests revealed that, for both the psychophysical and fMRI experiments, there was no significant difference between the 90° + 0° and 90° + 25° (the psychophysical experiment: *t*(24) = 0.446, *P* = 1.000, η_p_^2^ = 0.084; the fMRI experiment: *t*(19) = 2.166, *P* = 0.130, η_p_^2^ = 0.284) texture stimuli, and both were significantly higher than the 90° + 50° texture stimulus (the psychophysical experiment: 90° + 0° versus 90° + 50°: *t*(24) = 12.091, *P* < 0.001, η_p_^2^ = 2.738; 90° + 25° versus 90° + 50°: *t*(24) = 11.737, *P* < 0.001, η_p_^2^ = 2.712; the fMRI experiment: 90° + 0° versus 90° + 50°: *t*(19) = 11.257, *P* < 0.001, η_p_^2^ = 2.363; 90° + 25° versus 90° + 50°: *t*(19) = 10.449, *P* < 0.001, η_p_^2^ = 2.202). These results further indicate that our awareness manipulation was effective for both Visible and Invisible conditions. Error bars denote 1 SEM calculated across subjects and colored dots denote the data from each subject.

**Figure S2.**
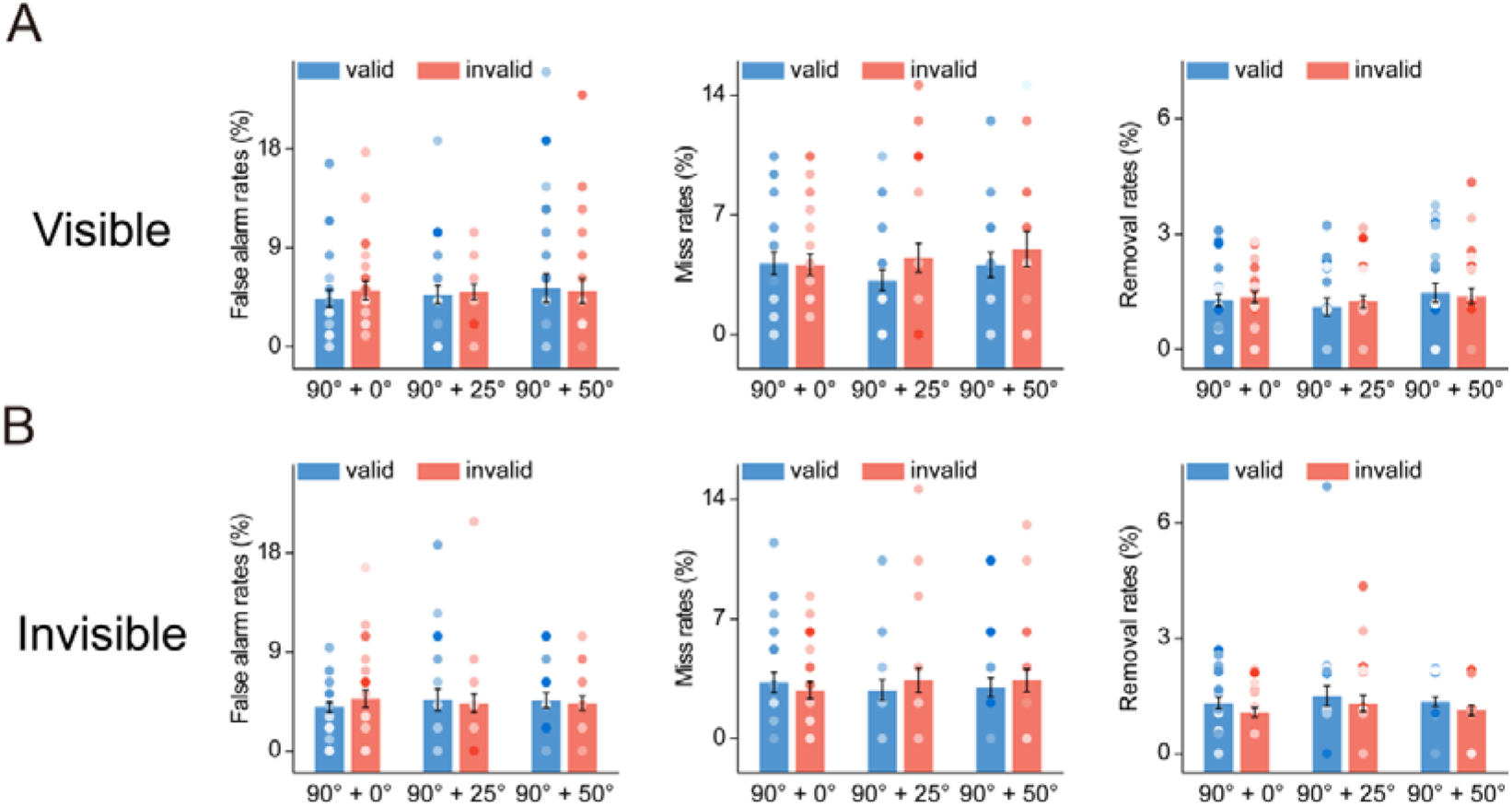
False alarm, Miss, and Removal Rates in Psychophysical Data. False alarm rates of 90° + 0°-valid cue, 90° + 0°-invalid cue, 90° + 25°-valid cue, 90° + 25°-invalid cue, 90° + 50°-valid cue, and 90° + 50°-invalid cue during Visible (**A**) and Invisible (**B**) conditions. Note that, in our study, subjects were asked to press one of two buttons as rapidly and correctly as possible to indicate the orientation of the ellipse probe (45° or 135°). Thus, for each condition, a rightward response to a 45° ellipse was (arbitrarily) considered to be a hit, a rightward response to a 135° ellipse was considered to be a false alarm, and a leftward response to a 45° ellipse was considered to be a miss. Error bars denote 1 SEM calculated across subjects and colored dots denote the data from each subject. Miss rates of those conditions during Visible (**A**) and Invisible (**B**) conditions. Error bars denote 1 SEM calculated across subjects and colored dots denote the data from each subject. Removal rates (i.e., correct reaction times shorter than 200 ms and beyond three standard deviations from the mean reaction time in each condition were removed) of those conditions during Visible (**A**) and Invisible (**B**) conditions. Error bars denote 1 SEM calculated across subjects and colored dots denote the data from each subject. There was no significant difference in false alarm rate, miss rate, or removal rate across conditions (all *P* > 0.05, η_p_^2^ < 0.151).

**Figure S3.**
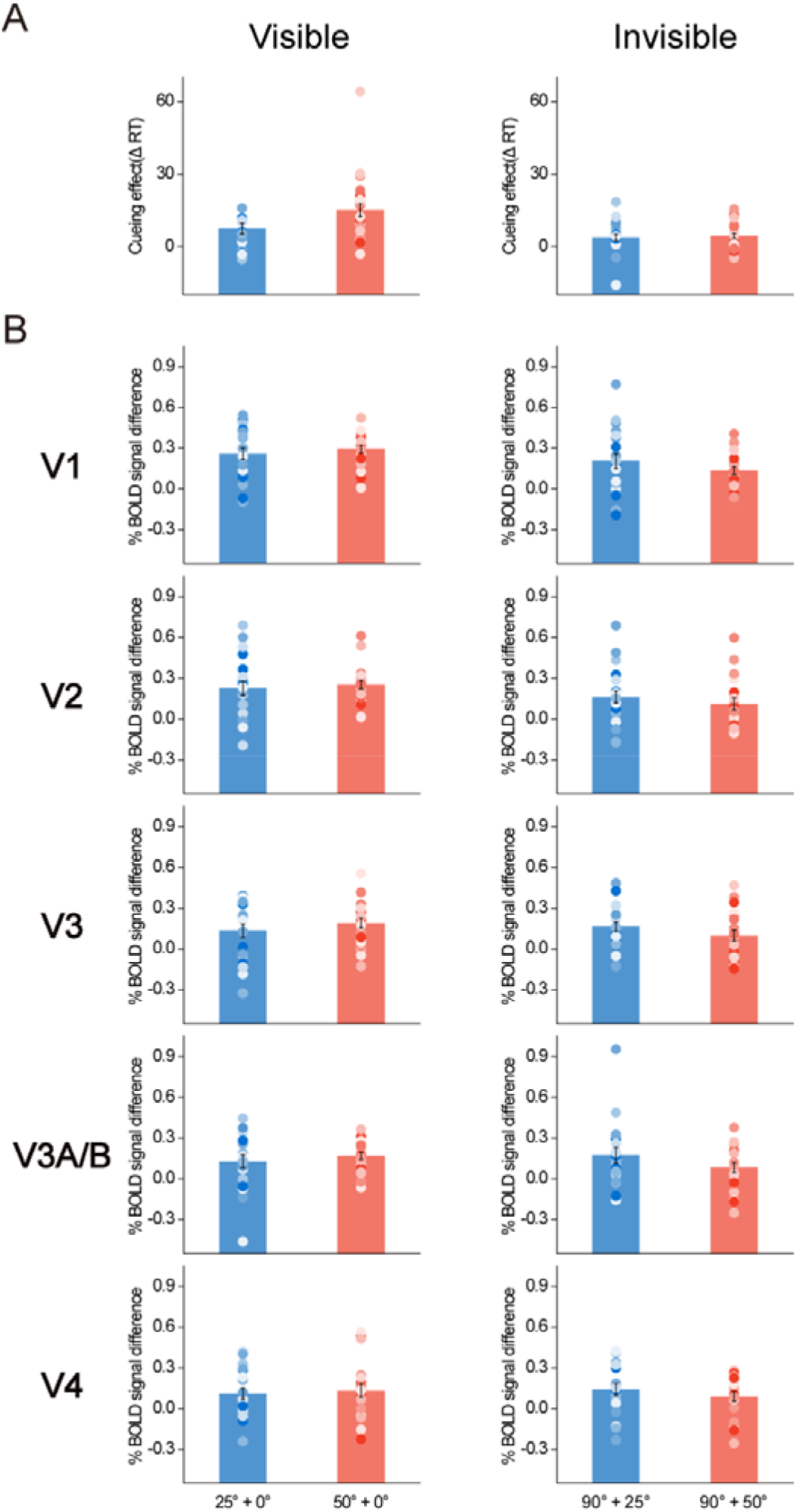
Cueing Effects and BOLD Signal Differences of 25°- and 50°-foregrounds. (**A**) The cueing effect of the 25°- and 50°-foregrounds for 25° + 0° and 50° + 0° texture stimuli, respectively, during the Visible (left) and Invisible (right) conditions. Each cueing effect was quantified as the difference between the reaction time of the probe task performance in the invalid cue condition and that in the valid cue condition. Error bars denote 1 SEM calculated across subjects and colored dots denote the data from each subject. (**B**) BOLD signal difference of the 25°- and 50°-foregrounds within 90° + 25° and 90° +50° texture stimuli, respectively, in SC and V1–V4 during the Visible (left) and Invisible (right) conditions. Error bars denote 1 SEM calculated across subjects and colored dots denote the data from each subject.

**Figure S4.**
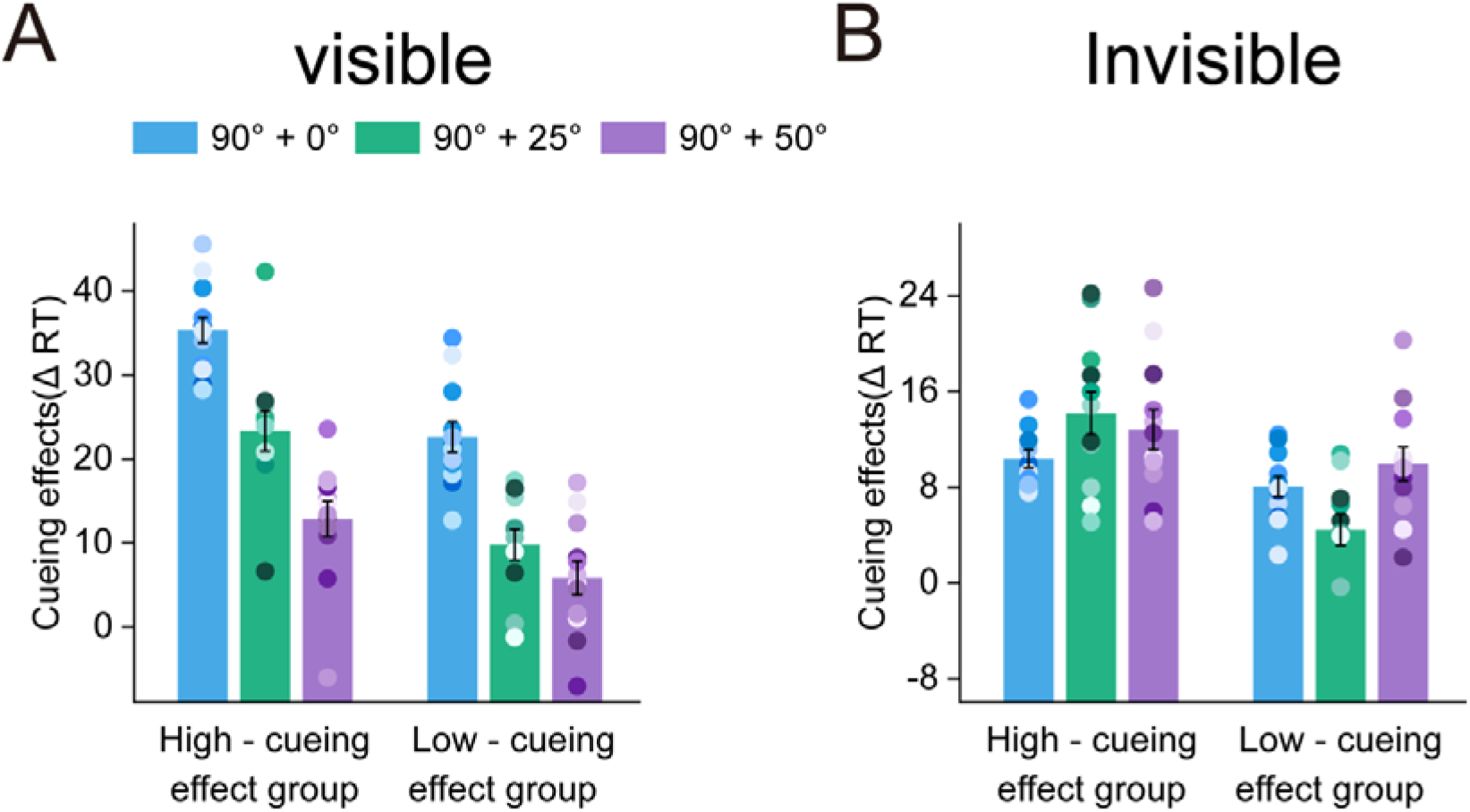
Cueing Effects of the High- and Low-cueing Effect Groups. Subjects who scored in the top and bottom 48% (i.e., *n* = 12) of the sample’s cueing effect distribution were assigned to High-cueing effect (left) and Low-cueing effect (right) groups, respectively, during both Visible (**A**) and Invisible (**B**) conditions. In both these two conditions, the cueing effects were submitted to a mixed ANOVA with group (Low- and High-cueing effect) as the between-subjects factor and texture stimulus (90° + 0°, 90° + 25°, and 90° + 50°) as the within-subjects factor. Results showed that, the interactions between the group and texture stimulus were not significant in both Visible (*F*(2,44) = 1.663, *P* = 0.207, η_p_^2^ = 0.07) and Invisible (*F*(2,44) = 3.597, *P* = 0.055, η_p_^2^ = 0.141) conditions, suggesting that the strength of cueing effect did not influence our conclusion that the saliency map was distributed as gradient or winner-take-all manner with or without awareness, respectively. Error bars denote 1 SEM calculated across subjects and colored dots denote the data from each subject.

**Figure S5.**
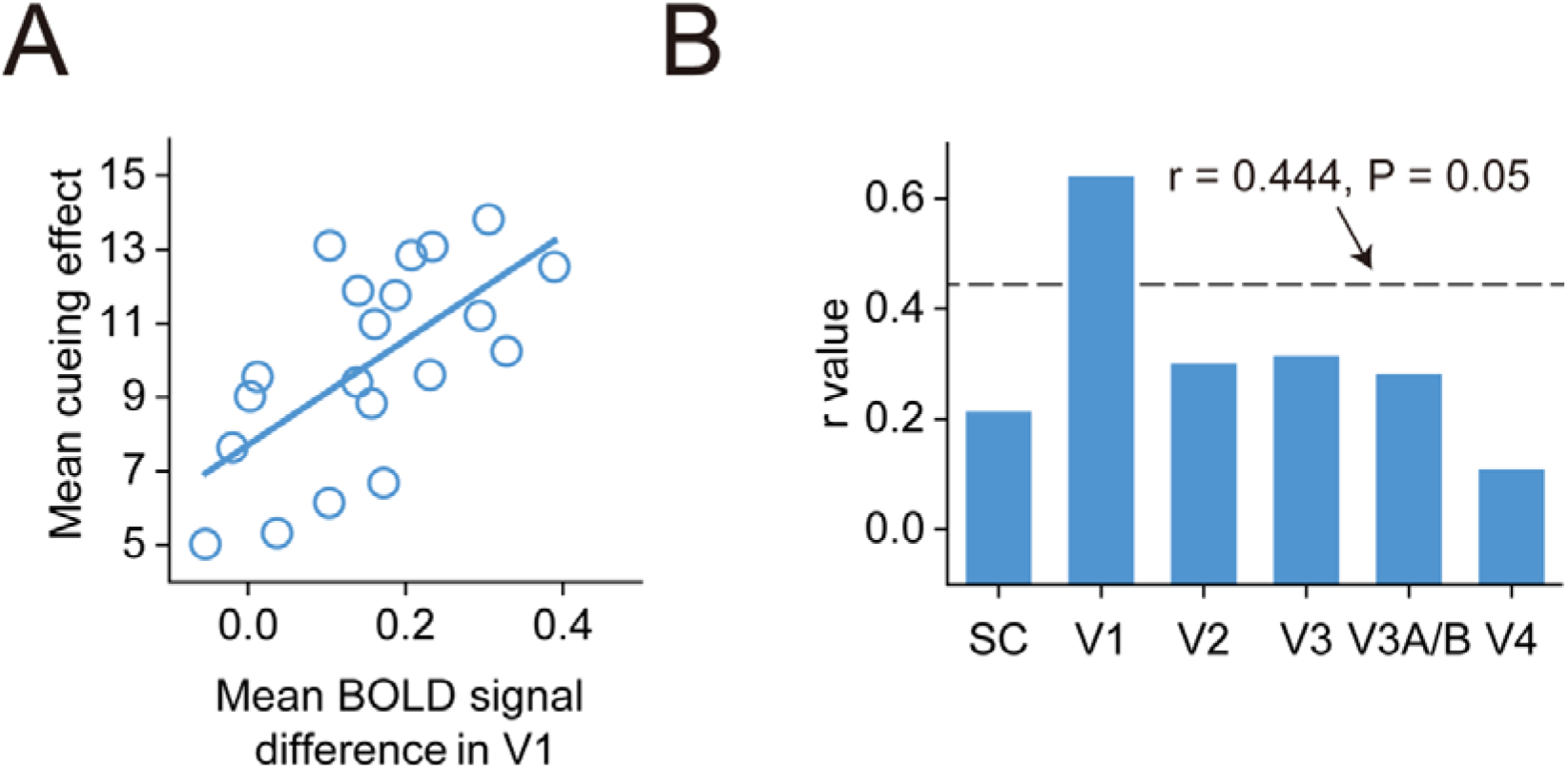
Correlations between the Mean Cueing Effect and the Mean BOLD Signal Difference for 90°-foreground of the Three Texture Stimuli. Correlations between the mean cueing effect and the mean BOLD signal difference for 90°-foreground of the three texture stimuli in V1 (**A**), and correlation coefficients (r values) between the mean cueing effect and the mean BOLD signal difference for 90°-foreground of the three texture stimuli in SC and V1–V4 (**B**), across individual subjects during the Invisible condition.

## Notes

https://osf.io/u7sbf/

### Competing Interest Statement

The authors have declared no competing interest.

## References

1. Allman J, Miezin F, McGuinness E. 1985. Stimulus specific responses from beyond the classical receptive field: neurophysiological mechanisms for local-global comparisons in visual neurons. Annu Rev Neurosci. 8: 407–430. https://doi.org/10.1146/annurev.ne.08.030185.002203 PMID: 3885829

2. Asplund CL, Todd JJ, Snyder AP, Marois R. 2010. A central role for the lateral prefrontal cortex in goal-directed and stimulus-driven attention. Nat Neurosci. 13: 507–512. https://doi.org/10.1038/nn.2509 PMID: 20208526

3. Baluch F, Itti L. 2011. Mechanisms of top-down attention. Trends Neurosci. 34: 210–224. https://doi.org/10.1016/j.tins.2011.02.003 PMID: 21439656

4. Basso MA, Wurtz RH. 2002. Neuronal activity in substantia nigra pars reticulata during target selection. J Neurosci. 22: 1883–1894. https://doi.org/10.1523/JNEUROSCI.22-05-01883.2002 PMID: 11880518

5. Bisley JW, Goldberg ME. 2010. Attention, intention, and priority in the parietal lobe. Annu Rev Neurosci. 33: 1–21. https://doi.org/10.1146/annurev-neuro-060909-152823 PMID: 20192813

6. Bogler C, Bode S, Haynes JD. 2011. Decoding successive computational stages of saliency processing. Curr Biol. 21: 1667–1671. https://doi.org/10.1016/j.cub.2011.08.039 PMID: 21962709

7. Boly M, Massimini M, Tsuchiya N, Postle BR, Koch C, Tononi G. 2017. Are the neural correlates of consciousness in the front or in the back of the cerebral cortex? Clinical and neuroimaging evidence. J Neurosci. 37: 9603–9613. https://doi.org/10.1523/JNEUROSCI.3218-16.2017 PMID: 28978697

8. Buffalo EA, Fries P, Landman R, Liang H, Desimone R. 2010. A backward progression of attentional effects in the ventral stream. Proc Natl Acad Sci USA. 107: 361–365. https://doi.org/10.1073/pnas.0907658106 PMID: 20007766

9. Burrows BE, Moore T. 2009. Influence and limitations of pop out in the selection of salient visual stimuli by area V4 neurons. J Neurosci. 29: 15169–15177. https://doi.org/10.1523/JNEUROSCI.3710-09.2009 PMID: 19955369

10. Buschman TJ, Miller EK. 2007. Top-down versus bottom-up control of attention in the prefrontal and posterior parietal cortices. Science 315: 1860–1862. https://doi.org/10.1126/science.1138071 PMID: 17395832

11. Cavanagh P, Alvarez GA. 2005. Tracking multiple targets with multifocal attention. Trends Cogn Sci. 9:349–354. https://doi.org/10.1016/j.tics.2005.05.009 PMID: 15953754

12. Chen C, Zhang X, Wang Y, Zhou T, Fang F. 2016. Neural activities in V1 create the bottom-up saliency map of natural scenes. Exp Brain Res. 234: 1769–1780. https://doi.org/10.1007/s00221-016-4583-y PMID: 26879771

13. Constantinidis C, Steinmetz MA. 2005. Posterior parietal cortex automatically encodes the location of salient stimuli. J Neurosci. 25:233–238. https://doi.org/10.1523/JNEUROSCI.3379-04.2005 PMID: 15634786

14. Corbetta M, Shulman GL. 2002. Control of goal-directed and stimulus driven attention in the brain. Nat Rev Neurosci. 3: 201–215. https://doi.org/10.1038/nrn755 PMID: 11994752

15. Corbetta M, Shulman GL. 2011. Spatial neglect and attention networks. Ann Rev Neurosci. 34: 569–599. https://doi.org/10.1146/annurev-neuro-061010-113731 PMID: 21692662

16. Del Cul A, Baillet S, Dehaene S. 2007. Brain dynamics underlying the nonlinear threshold for access to consciousness. PLoS Biol. 5: e260. https://doi.org/10.1371/journal.pbio.0050260 PMID: 17896866

17. Engel SA, Glover GH, Wandell BA. 1997. Retinotopic organization in human visual cortex and the spatial precision of functional MRI. Cereb Cortex. 7:181–192. https://doi.org/10.1093/cercor/7.2.181 PMID: 9087826

18. Fang F, Boyaci H, Kersten D, Murray SO. 2008. Attention-dependent representation of a size illusion in human V1. Curr Biol. 18:1707–1712. https://doi.org/10.1016/j.cub.2008.09.025 PMID: 18993076

19. Fecteau JH, Munoz DP. 2006. Salience, relevance, and firing: a priority map for target selection. Trends Cogn Sci. 10: 382–390. https://doi.org/10.1016/j.tics.2006.06.011 PMID: 16843702

20. Franconeri SL, Alvarez GA, Cavanagh P. 2013. Flexible cognitive resources: competitive content maps for attention and memory. Trends Cogn Sci. 17:134–141. https://doi.org/10.1016/j.tics.2013.01.010 PMID: 23428935

21. Friston KJ, Harrison L, Penny W. 2003. Dynamic causal modelling. NeuroImage 19: 1273–1302. https://doi.org/10.1016/s1053-8119(03)00202-7 PMID: 12948688

22. Friston KJ, Holmes AP, Worsley KJ, Poline JP, Frith CD, Frackowiak RS. 1994. Statistical parametric maps in functional imaging: a general linear approach. Hum. Brain Mapp. 2: 189–210. https://doi.org/10.1002/hbm.460020402

23. Geng JJ, Mangun GR. 2009. Anterior intraparietal sulcus is sensitive to bottom-up attention driven by stimulus salience. J Cogn Neurosci. 21: 1584–1601. https://doi.org/10.1162/jocn.2009.21103 PMID: 18752405

24. Genovese CR, Lazar NA, Nichols T. 2002. Thresholding of statistical maps in functional neuroimaging using the false discovery rate. NeuroImage 15: 870–878. https://doi.org/10.1006/nimg.2001.1037 PMID: 11906227

25. Gottlieb J. 2007. From thought to action: the parietal cortex as a bridge between perception, action, and cognition. Neuron 53:9–16. https://doi.org/10.1016/j.neuron.2006.12.009 PMID: 17196526

26. Gottlieb JP, Kusunoki M, Goldberg ME. 1998. The representation of visual salience in monkey parietal cortex. Nature 391: 481–484. https://doi.org/10.1038/35135 PMID: 9461214

27. Hegdé J, Felleman DJ. 2003. How selective are V1 cells for pop-out stimuli? J Neurosci. 23: 9968–9980. https://doi.org/10.1523/JNEUROSCI.23-31-09968.2003 PMID: 14602810

28. Hodsoll J, Mevorach C, Humphreys GW. 2009. Driven to less distraction: rTMS of the right parietal cortex reduces attentional capture in visual search. Cereb Cortex 19: 106–114. https://doi.org/10.1093/cercor/bhn070 PMID: 18515299

29. Huang L, Wang L, Shen W, Li M, Wang S, Wang X, Ungerleider LG, Zhang X. 2020. A source for awareness-dependent figure–ground segregation in human prefrontal cortex. Proc Natl Acad Sci USA. 117: 30836–30847. https://doi.org/10.1073/pnas.1922832117 PMID: 33199608

30. Itti L, Koch C. 2001. Computational modelling of visual attention. Nat Rev Neurosci. 2: 194–203. https://doi.org/10.1038/35058500 PMID: 11256080

31. Jonides J. 1981. Voluntary vs. automatic control over the mind’s eye’s movement. In Attention and Performance, Volume XI, M.I. Posner and O. Marin, eds. (Hillsdale, NJ: Lawrence Erlbaum Associates), 187–205.

32. Kanwisher N, Wojciulik E. 2000. Visual attention: insights from brain imaging. Nat Rev Neurosci. 1: 91–100. https://doi.org/10.1038/35039043 PMID: 11252779

33. Kastner S, De Weerd P, Desimone R, Ungerleider LG. 1998. Mechanisms of directed attention in the human extrastriate cortex as revealed by functional MRI. Science. 282:108–111. https://doi.org/10.1126/science.282.5386.108 PMID: 9756472

34. Kastner S, Nothdurft HC, Pigarev IN. 1997. Neuronal correlates of pop-out in cat striate cortex. Vis Res. 37: 371–376. https://doi.org/10.1016/s0042-6989(96)00184-8 PMID: 9156167

35. Kastner S, Ungerleider LG. 2000. Mechanisms of visual attention in the human cortex. Annu Rev Neurosci. 23: 315–341. https://doi.org/10.1146/annurev.neuro.23.1.315 PMID: 10845067

36. Katsuki F, Constantinidis C. 2012. Early involvement of prefrontal cortex in visual bottom-up attention. Nat Neurosci. 15: 1160–1166. https://doi.org/10.1038/nn.3164 PMID: 22820465

37. Klein RM. 2000. Inhibition of return. Trends Cogn Sci. 4: 138–147. https://doi.org/10.1016/S1364-6613(00)01452-2

38. Koch C, Ullman S. 1985. Shifts in selective visual attention: towards the underlying neural circuitry. Hum Neurobiol. 4: 219–227. PMID: 3836989

39. Koene AR, Zhaoping L. 2007. Feature-specific interactions in salience from combined feature contrasts: evidence for a bottom-up saliency map in V1. J Vis. 7: 1–14. https://doi.org/10.1167/7.7.6 PMID: 17685802

40. Kustov AA, Robinson DL. 1996. Shared neural control of attentional shifts and eye movements. Nature 384: 74–77. https://doi.org/10.1038/384074a0 PMID: 8900281

41. Liu T, Pestilli F, Carrasco M. 2005. Transient attention enhances perceptual performance and fMRI response in human visual cortex. Neuron. 45:469–477. https://doi.org/10.1016/j.neuron.2004.12.039 PMID: 15694332

42. Li Z. 1999. Contextual influences in V1 as a basis for pop out and asymmetry in visual search. Proc Natl Acad Sci USA. 96: 10530–10535. https://doi.org/10.1073/pnas.96.18.10530 PMID: 10468643

43. Li Z. 2002. A saliency map in primary visual cortex. Trends Cogn Sci. 6: 9–16. https://doi.org/10.1016/s1364-6613(00)01817-9 PMID: 11849610

44. Luck SJ, Chelazzi L, Hillyard SA, Desimone R. 1997. Neural mechanisms of spatial selective attention in areas V1, V2, and V4 of macaque visual cortex. J Neurophysiol. 77:24–42. https://doi.org/10.1152/jn.1997.77.1.24 PMID: 9120566

45. Mashour GA, Roelfsema P, Changeux JP, Dehaene S. 2020. Conscious processing and the global neuronal workspace hypothesis. Neuron 105: 776–798. https://doi.org/10.1016/j.neuron.2020.01.026 PMID: 32135090

46. Mazer JA, Gallant JL. 2003. Goal-related activity in V4 during free viewing visual search. Evidence for a ventral stream visual salience map. Neuron 40: 1241–1250. https://doi.org/10.1016/s0896-6273(03)00764-5 PMID: 14687556

47. Melloni L, van Leeuwen S, Alink A, Müller NG. 2012. Interaction between bottom-up saliency and top-down control: how saliency maps are created in the human brain. Cereb Cortex. 22:2943–2952. https://doi.org/10.1093/cercor/bhr384 PMID: 22250291

48. Mevorach C, Hodsoll J, Allen H, Shalev L, Humphreys G. 2010. Ignoring the elephant in the room: a neural circuit to downregulate salience. J Neurosci. 30: 6072–6079. https://doi.org/10.1523/JNEUROSCI.0241-10.2010 PMID: 20427665

49. Mevorach C, Humphreys GW, Shalev L. 2006. Opposite biases in salience-based selection for the left and right posterior parietal cortex. Nat Neurosci. 9: 740–742. https://doi.org/10.1038/nn1709 PMID: 16699505

50. Mevorach C, Humphreys GW, Shalev L. 2009. Reflexive and preparatory selection and suppression of salient information in the right and left posterior parietal cortex. J Cogn Neurosci. 21: 1204–1214. https://doi.org/10.1162/jocn.2009.21088 PMID: 18752407

51. Nakayama K, Mackeben M. 1989. Sustained and transient components of focal visual attention. Vision Res. 29: 1631–1647. https://doi.org/10.1016/0042-6989(89)90144-2 PMID: 2635486

52. Nothdurft HC, Gallant JL, Van Essen DC. 1999. Response modulation by texture surround in primate area V1: correlates of ‘‘pop out’’ under anesthesia. Vis Neurosci. 16: 15–34. https://doi.org/10.1017/s0952523899156189 PMID: 10022475

53. Payne HE, Allen HA. 2011. Active ignoring in early visual cortex. J Cogn Neurosci. 23: 2046–2058. https://doi.org/10.1162/jocn.2010.21562 PMID: 20807054

54. Penny WD, Stephan KE, Mechelli A, Friston KJ. 2004. Modelling functional integration: a comparison of structural equation and dynamic causal models. NeuroImage. 23:S264–S274. https://doi.org/10.1016/j.neuroimage.2004.03.026 PMID: 15219588

55. Posner MI, Snyder CRR, Davidson BJ. 1980. Attention and the detection of signals. J Exp Psychol. 109: 160–174. PMID: 7381367

56. Ruff CC, Driver J. 2006. Attentional preparation for a lateralized visual distractor: Behavioral and fMRI evidence. J Cogn Neurosci. 18: 522–538. https://doi.org/10.1162/jocn.2006.18.4.522 PMID: 16768358

57. Serences JT, Shomstein S, Leber AB, Golay X, Egeth HE, Yantis, S. 2005. Coordination of voluntary and stimulus-driven attentional control in human cortex. Psychol Sci. 16: 114–122. https://doi.org/10.1111/j.0956-7976.2005.00791.x PMID: 15686577

58. Serences JT, Yantis S. 2006. Selective visual attention and perceptual coherence. Trends Cogn Sci. 10: 38–45. https://doi.org/10.1016/j.tics.2005.11.008 PMID: 16318922

59. Serences JT, Yantis S. 2007. Spatially selective representations of voluntary and stimulus-driven attentional priority in human occipital, parietal, and frontal cortex. Cereb Cortex. 17: 284–293. https://doi.org/10.1093/cercor/bhj146 PMID: 16514108

60. Sereno MI, Dale AM, Reppas JB, Kwong KK, Belliveau JW, Brady TJ, Rosen BR, Tootell RBH. 1995. Borders of multiple visual areas in humans revealed by functional magnetic resonance imaging. Science. 268:889–893. https://doi.org/10.1126/science.7754376 PMID: 7754376

61. Shipp S. 2004. The brain circuitry of attention. Trends Cogn Sci. 8: 223–230. https://doi.org/10.1016/j.tics.2004.03.004 PMID: 15120681

62. Shomstein S. 2012. Cognitive functions of the posterior parietal cortex: top-down and bottom-up attentional control. Front Integr Neurosci. 6: 38. https://doi.org/10.3389/fnint.2012.00038 PMID: 22783174

63. Smith AM, Lewis BK, Ruttimann UE, Ye FQ, Sinnwell TM, Yang Y, Duyn JH, Frank JA. 1999. Investigation of low frequency drift in fMRI signal. Neuroimage. 9:526–533. https://doi.org/10.1006/nimg.1999.0435 PMID: 10329292

64. Squire RF, Noudoost B, Schafer RJ, Moore T. 2013. Prefrontal contributions to visual selective attention. Ann Rev Neurosci. 36: 451–466. https://doi.org/10.1146/annurev-neuro-062111-150439 PMID: 23841841

65. Talairach J, Tournoux P. 1998. Co-Planar Stereotaxic Atlas of the Human Brain: 3-Dimensional Proportional System: an Approach to Cerebral Imaging (New York: Thieme).

66. Thompson KG, Bichot NP. 2005. A visual salience map in the primate frontal eye field. Prog Brain Res. 147: 251–262. https://doi.org/10.1016/S0079-6123(04)47019-8 PMID: 15581711

67. White BJ, Berg DJ, Kan JY, Marino RA, Itti L, Munoz DP. 2017a. Superior colliculus neurons encode a visual saliency map during free viewing of natural dynamic video. Nat Commun. 8: 1–9. https://doi.org/10.1038/ncomms14263 PMID: 28117340

68. White BJ, Kan JY, Levy R, Itti L, Munoz DP. 2017b. Superior colliculus encodes visual saliency before the primary visual cortex. Proc Natl Acad Sci USA. 114: 9451–9456. https://doi.org/10.1073/pnas.1701003114 PMID: 28808026

69. Wolfe JM. 1994. Visual search in continuous, naturalistic stimuli. Vision Res. 34: 1187–1195. https://doi.org/10.1016/0042-6989(94)90300-x PMID: 8184562

70. Yan Y, Zhaoping L, Li W. 2018. Bottom-up saliency and top-down learning in the primary visual cortex of monkeys. Proc Natl Acad Sci USA. 115: 10499–10504. https://doi.org/10.1073/pnas.1803854115 PMID: 30254154

71. Zhaoping L. 2008. Attention capture by eye of origin singletons even without awareness—a hallmark of a bottom-up saliency map in the primary visual cortex. J Vis. 8: 1–18. https://doi.org/10.1167/8.5.1 PMID: 18842072

72. Zhaoping L, May KA. 2007. Psychophysical tests of the hypothesis of a bottom-up saliency map in primary visual cortex. PLoS Comput Biol. 3: e62. https://doi.org/10.1371/journal.pcbi.0030062 PMID: 17411335

73. Zhaoping L, Zhe L. 2015. Primary visual cortex as a saliency map: a parameter-free prediction its test by behavioral data. PLoS Comput Biol. 11: e1004375. https://doi.org/10.1371/journal.pcbi.1004375 PMID: 26441341

74. Zhang X, Japee S, Safiullah Z, Mlynaryk N, Ungerleider LG. 2016. A normalization framework for emotional attention. PLoS Biol. 14: e1002578. https://doi.org/10.1371/journal.pbio.1002578 PMID: 27870851

75. Zhang X, Mlynaryk N, Ahmed S, Japee S, Ungerleider LG. 2018. The role of inferior frontal junction in controlling the spatially global effect of feature-based attention in human visual areas. PLoS Biol. 16: e2005399. https://doi.org/10.1371/journal.pbio.2005399 PMID: 29939981

76. Zhang X, Qiu J, Zhang Y, Han S, Fang F. 2014. Misbinding of color and motion in human visual cortex. Curr Biol. 24:1354–1360. https://doi.org/10.1016/j.cub.2014.04.045 PMID: 24856212

77. Zhang X, Zhaoping L, Zhou T, Fang F. 2012. Neural activities in V1 create a bottom-up saliency map. Neuron 73: 183–192. https://doi.org/10.1016/j.neuron.2011.10.035 PMID: 22243756

